# A recombinant heterotrimeric mini-procollagen I reveals new functional and structural roles for the α2(I) chain

**DOI:** 10.64898/2026.10.03.756409

**Authors:** Oskar Lipinski, Cindy Dieryckx, Natacha Mariano, Prisicillia Lagoutte, Hok Sau Kwong, Sandrine Vadon-Le Goff, Loïc Carrique, Catherine Moali

## Abstract

Collagen I provides mechanical stability to organs and plays a critical role in several physiological processes such as wound healing and bone remodeling. Its ability to form fibrils depends on the proteolytic conversion of soluble procollagen I into mature protomers. However, the mechanistic aspects of this critical step have been difficult to analyze due to the lack of recombinant procollagen I suitable for structural and biochemical analyses. Here, we describe the successful production and purification of a heterotrimeric mini-procollagen I [α1(I)_2_α2(I)] together with its homotrimeric counterpart [α1(I)_3_] in HEK293-F cells. These tools were used to probe the specific role of the α2(I) chain both in procollagen I assembly and proteolytic maturation. While the presence of α2(I) in the heterotrimer does not strengthen trimer interfaces on its own, as revealed by cryo-electron microscopy, it provides a clear functional advantage to the procollagen substrate that is cleaved more efficiently by bone morphogenetic protein 1 (BMP-1) than the homotrimer. Furthermore, we demonstrate that the first cleavage event in the heterotrimer is at the level of the α2(I) chain, through a mechanism involving its C-telopeptide and a specific PRS motif close to the cleavage site. The preferential cleavage of α2(I) by BMP-1 can be partially corrected in the presence of PCPE-1 (procollagen C-proteinase enhancer 1) but the efficacy of the correction depends on its position on the procollagen trimer. Finally, we solved the high-resolution structure of the complex between the heterotrimer and PCPE-1 and proposed a new mechanism to explain the enhancing activity of PCPE-1.

## Introduction

Fibrillar collagens are composed of a central triple helical domain, made of more than 300 Gly-Xaa-Yaa repeats, flanked by N- and C-telopeptide regions. They are synthesized as procollagens which also contain additional domains at both extremities, called N- and C-propeptides (1, 2). These domains maintain procollagens in soluble form until they are removed proteolytically by procollagen N- and C-proteinases (PNPs and PCPs), a critical step to trigger fibril formation.

Among fibrillar collagens, type I collagen is the most abundant protein in several human connective tissues including skin, cornea, tendon, ligament and bone. While (pro)collagen I is predominantly found in tissues as a heterotrimer consisting of two α1(I) chains and one α2(I) chain, increased levels of homotrimeric (pro)collagen I, made of three α1(I) chains, are also observed during development (3) and in several disease contexts. A well-described example is the tumor microenvironment where homotrimeric procollagen I is produced by cancer cells upon epigenetic repression of the *COL1A2* gene encoding the α2(I) chain and contributes to tumor progression (4–6). Homotrimers can also be substantially over-represented in fibrotic disorders such as cirrhosis (7) or Dupuytren’s disease (8) and in genetic diseases such as osteogenesis imperfecta (9) or Ehlers-Danlos syndrome (10).

Collagen I homotrimers were previously reported to behave differently from heterotrimers in terms of thermal stability (11), ability to form fibrils (9) or susceptibility to degradation by matrix metalloproteinases (MMPs) (12). However, to our knowledge, the efficiencies of the proteolytic conversion of homo- and hetero-trimeric procollagens into mature collagens by PNPs and PCPs have never been thoroughly compared. Also, it is presently unclear why the formation of heterotrimers is so much favored over the formation of homotrimers *in vivo*. Crystal structures of homotrimeric C-propeptides suggest that trimer assembly relies on specific interactions between α chains which are further stabilized by calcium ions and inter-chain disulfide bonds, as also confirmed biochemically (2, 13, 14). The fact that heterotrimeric procollagen I can be formed despite the lack of one disulfide bond at one of the α1-α2 interfaces is therefore puzzling and the mechanisms governing heterotrimerization remain elusive. In addition to chain recognition at the level of C-propeptides, it was also recently demonstrated that the propagation of triple helices with appropriate stoichiometries is further regulated by the nature of the residues located at the C-terminal end of the collagenous domain (15).

A better understanding of these assembly and maturation steps would give important insights into how tissue-specific properties of collagen fibrils are established and maintained. Moreover, it would also help to design appropriate therapeutic strategies to control deficient or excessive collagen I deposition, for example in impaired wound healing, fibrosis or genetic diseases. However, so far, this type of functional and structural studies has been hampered by the difficulty to obtain large amounts of pure heterotrimeric procollagen I.

Native procollagen I can be successfully enriched in milligram amounts from the supernatant of skin or tendon fibroblasts cultivated in the presence of ascorbic acid (16, 17) but partial proteolysis by endogenous PNPs and PCPs often occurs, making separation of processed and unprocessed forms challenging. Recombinant expression of full-length procollagen I was also achieved with very high yields in yeast *Pichia pastoris* or tobacco plants through the co-expression of prolyl-4-hydroxylase to ensure proper proline 4-hydroxylation (18–20). In these systems, procollagens were not secreted, yet allowing the successful extraction and purification of both procollagen I and pepsin-digested collagen I (19–21). However, these proteins miss some important post-translational modifications such as proline 3-hydroxylation or lysine hydroxylation and, although the α1 and α2 chains appeared to be present with the expected stoichiometry of 2, it cannot be excluded that a substantial amount of homotrimer was also co-purified. Other common production systems for procollagens are mammalian cells such as CHO, HT1080 or HEK293 cells (15, 22–25) which have endogenous prolyl- and lysyl-hydroxylase activity. The use of these systems still poses several challenges related to the separation of the different types of trimers occurring when more than one α chain is transfected, and to the co-purification of several collagen-bound contaminants.

Based on the numerous pitfalls of the production and purification of full-length procollagens, we have developed recombinant mini-procollagens which correspond to truncated versions of human procollagens and can be tailored for various applications. Especially, we previously designed and characterized several mini-procollagens derived from procollagen III which, in addition to the C-propeptide, included N-terminal extensions of variable lengths. These constructs proved extremely useful for the characterization of metalloproteinases with procollagen C-proteinase activity, namely BTPs (BMP-1 (bone morphogenetic protein-1)/Tolloid-like Proteinases) (23, 26), meprins α and β (21, 27), ADAMTS (A Disintegrin And Metalloprotease with ThromboSpondin motifs) -2 and -14 (28), but also of the regulation of these maturation events by PCPE-1 (procollagen C-proteinase enhancer protein-1) and PCPE-2 (26, 27, 29). For example, successive truncations of a mini-procollagen III allowed us to prove that BMP-1 can cleave a minimal protein composed of the C-propeptide of human procollagen III and a short N-terminal tail (23) and that PCPE-1 enhances procollagen processing by BMP-1 through C-propeptide binding (23, 26). Short mini-procollagen III variants also proved very useful for structural analysis by SAXS and X-ray crystallography (13, 30, 31).

In the present work, we have built on this previous experience of mini-procollagen design to address the challenge of producing a heterotrimeric human mini-procollagen I. The production of this protein, together with the corresponding homotrimer, and their extensive characterizations are described below. Using these tools, we solved the structure of heterotrimeric mini-procollagen I in complex with PCPE-1 and identified the first chain that is cleaved by BMP-1 upon PCPE-1 binding. We also evidenced that the α2(I) chain provides a functional advantage to the heterotrimer through an increased cleavage rate by BMP-1 and a higher affinity for PCPE-1. However, it does not significantly promote the stabilization of the trimer interfaces on its own and additional factors such as the presence of calcium seem to play a role. Finally, our data suggest a new model for the mechanism of action of the three-partner complex involving procollagen I, BMP-1 and PCPE-1.

## Results

### Production and purification of heterotrimeric and homotrimeric mini-procollagens

Previous work has shown that it is possible to recombinantly express the heterotrimeric human C-propeptide I (CPI) in HEK293 cells and to separate this protein from α1(I)_3_ homotrimers through the introduction of a specific tag on the α2(I) chain (2). Importantly, the cysteine pattern of the α2(I) chain precludes the stable homotrimerization of this chain and dictates its association with two α1(I) chains to form α1(I)_2_α2(I) heterotrimers (14). We decided to use a similar strategy to extend the length of the α1(I) and α2(I) chains in the N-terminal direction, including the C-telopeptides and C-terminal ends of triple helical domains found in native human sequences, and additional GPP repeats to lock the N-terminal end of the triple helix (Fig. 1A; Fig. S1). Considering that the GGG sequence located at the end of the collagenous domain of the α2(I) chain is more likely to belong to the C-telopeptide than to the triple helix (15), the expected length of the triple helical domain is 35 triplets. A Twin-Strep-tag® (32) was also specifically incorporated at the N-terminus of the α2(I) chain and the N-glycosylation sites in the C-propeptide sequences were removed through mutagenesis to facilitate structural characterization (Fig. S1) (2, 13, 33). For protein production, we tested two culture systems allowing the transient transfection of either adherent (HEK293-T) or suspension (HEK293-F) cells. As expression levels in the two systems appeared to be similar (Fig. S2A), we chose to work with the HEK293-F cells that offer the advantage of growing in defined media, leading to less contaminants and better reproducibility.

**Figure 1:**
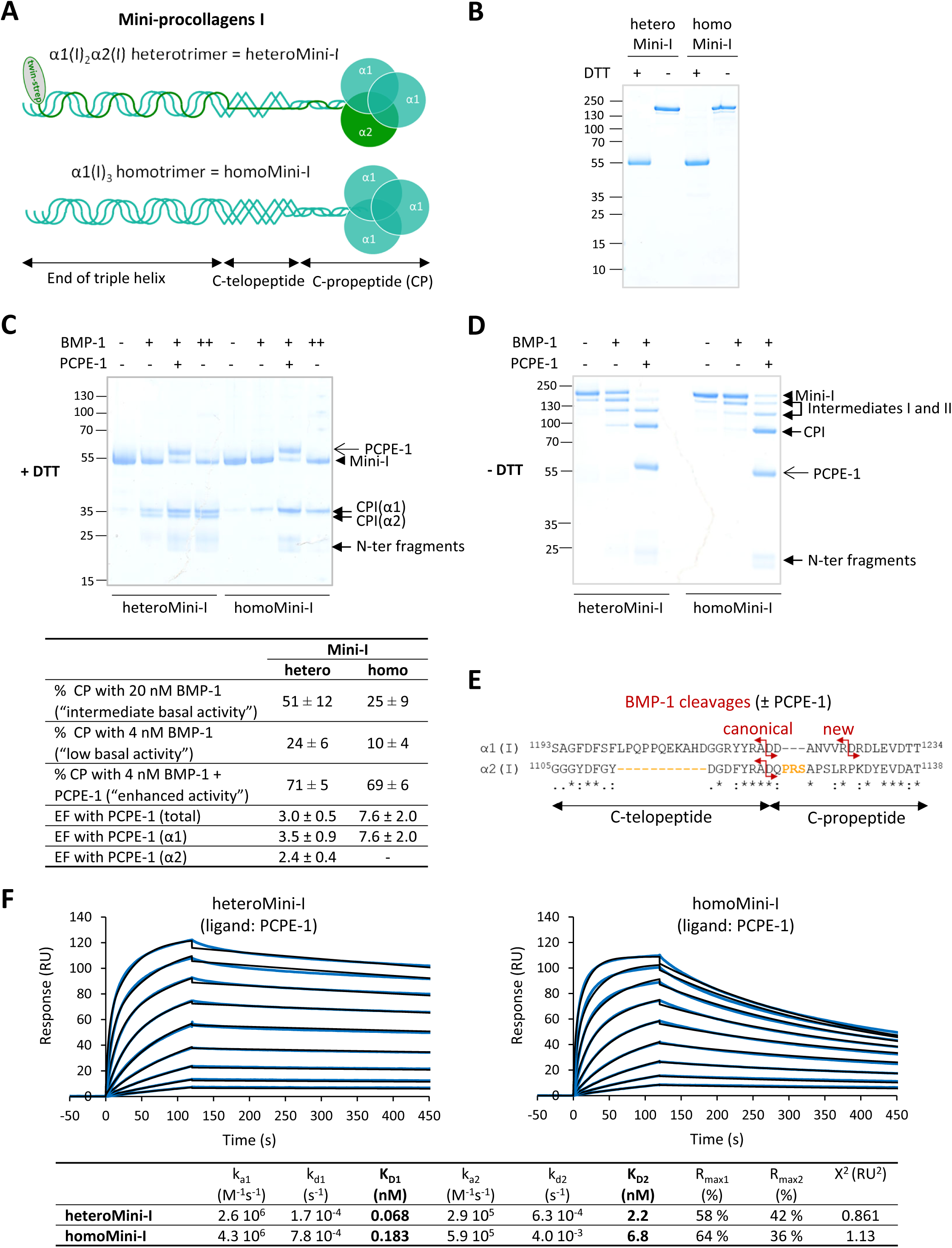
Design, production and functional characterization of mini-procollagens I. **(A)** HeteroMini-I and homoMini-I are composed of native C-propeptides (CPs), native C-telopeptides and truncated triple helices as described in Fig. S1. The α2(I) chain also contains a Twin-Strep-tag® at the N-terminus. **(B)** Trimer formation in heteroMini-I and homoMini-I is evidenced by a shift in molecular weight upon DTT reduction (4-12 % acrylamide gradient gel, InstantBlue staining). **(C)** SDS-PAGE analysis of the cleavage of heteroMini-I and homoMini-I (450 nM) incubated with BMP-1 (4 nM, indicated by “+”; or 20 nM, indicated by “++”), in the absence or presence of PCPE-1 (450 nM) for 30 min. The gel (4–12 % acrylamide gradient) was obtained in reducing conditions after Instant Blue staining and is representative of n = 4 (homotrimer) or n = 6 (heterotrimer) independent experiments which were quantified to yield the cleavage efficiencies and enhancement factors (EF) (calculated as described in Materials and Methods, in the presence of 4 nM BMP-1 and 450 nM PCPE-1). Means ± SD. **(D)** Same as in (C) except that the time was 45 min and the gel was run in non-reducing conditions. **(E)** Alignment of the C-telopeptides and the N-terminal regions of the CPs of the α1(I) and α2(I) chains obtained with ClustalO, highlighting in red the BMP-1 cleavage sites (see Table 1) and in orange the distinct features of the α2(I) chain, including a deletion in the C-telopeptide and a 3 amino-acid insertion in the CP. **(F)** Interaction of heteroMini-I and homoMini-I with immobilized PCPE-1 (140 RU). Increasing concentrations of heteroMini-I and homoMini-I were injected (0.39-100 nM, prepared as serial two-fold dilutions, with one duplicate injection of the 6.25 nM concentration; blue curves). The best fit of the experimental curves obtained with the heterogenous ligand model is also shown with the corresponding kinetic and dissociation constants (black curves).

**Table 1:** Cleavage site identification by ATOMS. HeteroMini-I and homoMini-I (950 nM) were incubated either with 24 nM BMP-1 or with 46 nM BMP-1 and 950 nM PCPE-1 for 1 h and, after TMT-labeling, their N-terminal peptides were compared to the control condition (without BMP-1 and PCPE-1). ATOMS ratios were calculated as the averages of all the ratios obtained for peptides defining cleavage sites (↓) or mature N-termini (after signal peptide removal) of the respective mini-procollagens. Cleavage site positions are indicated as in native human proteins, identified by their Uniprot ID, and previously described cleavage sites are marked by an asterisk (*). The working enzyme used to digest proteins could be trypsin or chymotrypsin.

| Reference protein | Working enzyme | Mature N-terminus or cleavage site | Protease/control ratio in heteroMini-I (number of peptide identifications) |  | Protease/control ratio in homoMini-I (number of peptide identifications) |  |
| --- | --- | --- | --- | --- | --- | --- |
|  |  |  | BMP-1 | BMP-1 + PCPE-1 | BMP-1 | BMP-1 + PCPE-1 |
| Collagen I (α1) P02452 | chymotrypsin / trypsin | Mature N-terminus in α1(I) | 0.8 (11) | 0.5 (11) | 0.9 (55) | 0.9 (55) |
|  | chymotrypsin / trypsin | YYRA↓<sup>1219</sup>DDAN* | 4.4 (8) | 6.9 (8) | 82 (6) | 97 (6) |
|  | chymotrypsin / trypsin | NVVR↓<sup>1226</sup>DRDL | 3.1 (7) | 10.4 (7) | 0.9 (7) | 3.2 (7) |
| Collagen I (α2) P08123 | chymotrypsin / trypsin | Mature N-terminus in α2(I) | 0.6 (3) | 0.2 (3) | - | - |
|  | chymotrypsin | FYRA↓<sup>1120</sup>DQPR* | 6.7 (7) | 5.1 (7) | - | - |

Purification was then achieved in two steps, affinity purification through the tag, where homotrimers consisting of α1(I) chains only were eliminated in the flow-through (Fig. S2B), and size exclusion chromatography (SEC). Based on the quantification of the α1(I) signal in flow-through and elution fractions, it was estimated that the proportions of homo- and hetero-trimers were 12 % and 88 %, respectively, in our production system (Fig. S2C). We also transfected the α1(I) construct alone in HEK293-F cells to obtain the corresponding homotrimeric mini-procollagen I which was purified through three chromatographic steps, in the absence of any tag (see Materials and Methods). Both proteins, called heteroMini-I and homoMini-I, trimerize as indicated by the shift upon reduction by SDS-PAGE and are more than 95 % pure (Fig. 1B) with less than 1 % contamination by endogenous PCPE-1 (Fig. S2D) and typical yields after purification in the range 3-6 mg/L.

During our preliminary characterizations, we also observed that the triple helix of both proteins was well-folded and resistant to pepsin digestion (Fig. S2E). As calcium is known to play a major role in stabilizing C-propeptide structures (2, 13, 14, 34), we next analyzed the impact of calcium depletion on hetero- and homo-trimer stability using NanoDSF (Differential Scanning Fluorometry) (Fig. S2F). We observed that, in the presence of 5 mM calcium, both types of trimers showed two distinct unfolding events with the second transition being clearly shifted towards higher temperatures for homoMini-I compared to heteroMini-I, indicating a markedly higher thermal stability of the corresponding subdomain (melting temperature of 86°C compared to 71°C). This may be attributed to one additional inter-chain disulfide bridge in homoMini-I. Upon chelation of calcium ions with 20 mM EDTA, the high-temperature transition disappeared, while the lower-temperature transition was shifted towards lower temperatures. Thus, the thermal stability of both heteroMini-I and homoMini-I was clearly linked to the presence of calcium.

### Cleavage of mini-procollagens by BMP-1 and interaction with PCPE-1

The next step was to validate the ability of BMP-1, the major PCP, to efficiently cleave heteroMini-I and homoMini-I and of PCPE-1 to enhance these cleavages. To this end, we incubated the mini-procollagens (450 nM) with BMP-1 (4 or 20 nM) at 37 °C and analyzed the cleavage products by SDS-PAGE, in both reducing (Fig. 1C) and non-reducing conditions (Fig. 1D). Both recombinant mini-procollagens could be cleaved by BMP-1 and these cleavages were stimulated by PCPE-1 (here in a 1:1 molar ratio to mini-procollagens), as expected. Based on the quantification of the amounts of C-propeptides (CPs), observed in the presence of DTT after cleavage at the two BMP-1 concentrations (no PCPE-1), we found that heteroMini-I was cleaved at least twofold more extensively than homoMini-I (Fig. 1C). This is a remarkable difference considering that the two proteins only differ at the level of one chain. After BMP-1 cleavage, the two bands corresponding to the CPs of the α1(I) and α2(I) chains of heteroMini-I were clearly resolved by SDS-PAGE in reducing conditions and confirmed that the two chains were present and covalently linked. Finally, the cleavage pattern in non-reducing conditions (Fig. 1D) indicated that processing of both mini-procollagens occurs through distinct intermediate products, corresponding to the successive cleavages of the three chains before the CP can be fully released.

In the presence of PCPE-1, the difference in cleavage efficiency between heteroMini-I and homoMini-I was completely abolished (Fig. 1C), suggesting that PCPE-1 can normalize the lower activity of BMP-1 observed with homoMini-I. This was confirmed by the determination of the enhancement factor (EF), calculated in conditions of low BMP-1 basal activity (4 nM BMP-1, 30 min) as the ratio of the % CP cleavage with PCPE-1 over the % CP cleavage without PCPE-1, which was 2.5 higher for homoMini-I than for heteroMini-I (Fig. 1C). Overall, our data indicate that PCPE-1 enhances the cleavage of the homotrimer more efficiently than the cleavage of the heterotrimer.

We then decided to confirm the previously described cleavage sites for BMP-1 in procollagen I (35). We used a mass-spectrometry-based approach called ATOMS (Amino-Terminal Oriented Mass spectrometry of Substrates) which relies on the specific isotopic labeling of N-termini, either naturally present in proteins or generated by proteolysis (36, 37). The mini-procollagens were incubated alone (control condition), with BMP-1 (protease condition) or with BMP-1 and PCPE-1 (enhanced protease condition) and cleavage sites were revealed by N-terminally-labeled peptides with isotopic ratios (protease/control or enhanced protease/control) superior to 3. Interestingly, using this approach, we confirmed the previously described cleavage sites in α1(I) and α2(I) chains (marked with an asterisk in Table 1), which both have an aspartate in P1’ position in good agreement with the known substrate specificity of BMP-1 (38). However, we also found one additional cleavage in the α1 chains of both substrates, suggesting that previous information was not complete. This new cleavage occurred 7 residues C-terminal to the previously described site and also had an aspartate in the P1’ position (Fig. 1E). Importantly, the same cleavage sites were identified in the absence and presence of PCPE-1, indicating that PCPE-1 does not change the cleavage specificity of BMP-1. The second cleavage site may also explain why the N-terminal fragments released from homoMini-I appear as two distinct bands by SDS-PAGE (Fig. 1C-D).

The enhancing activity of PCPE-1 was previously found to depend on its tight interaction with C-propeptides, as demonstrated for native CPI and for recombinant CPIII (26, 30, 39). Therefore, we also analyzed the binding of mini-procollagens I to immobilized PCPE-1 by Surface Plasmon Resonance (SPR) (Fig. 1F). As expected, both hetero- and homo-trimeric mini-procollagens could interact with PCPE-1. However, the binding profiles differed substantially and heteroMini-I formed a more stable complex with PCPE-1, as demonstrated by its slower dissociation. These findings were confirmed by successive injections of serial dilutions of the mini-procollagens and curve fitting with the heterogenous ligand model which was previously found to give the best results with mini-procollagen III (26, 40). This model yielded two dissociation constants for each mini-procollagen which were all below 6.8 nM (Fig. 1F), indicating that both heteroMini-I and homoMini-I were very strong ligands of PCPE-1. This analysis also confirmed that the biggest difference between the two trimers was at the level of the off-rates which were approximately 5-fold higher for homoMini-I, leading to slightly higher K_D_s. Given that the cleavage of homoMini-I is more efficiently enhanced by PCPE-1 than that of heteroMini-I (Fig. 1C), it seems that there is no direct positive correlation between the binding affinity for PCPE-1 and the level of stimulation.

All these characterizations validated the use of mini-procollagens as model substrates to analyze procollagen C-proteinase activity and to gain mechanistic insights into the regulation of the procollagen maturation complexes by PCPE-1 or other regulatory proteins.

### Preferential cleavage of α2(I) by BMP-1

While performing cleavage experiments, we noticed that the ratio between the α1 and α2 chains in the CP products of heteroMini-I (expected to be 2:1) was changing with the progress of the cleavage reaction. It was increasing from 1 at the earliest tested time point/lowest tested protease concentration to up to 1.8 when the cleavage was almost complete (Fig. 2A). Together with the comparatively slower cleavage of homoMini-I described above (Fig. 1C), this suggested that the cleavage of α1(I) by BMP-1 might be less efficient than the cleavage of α2(I). This observation prompted us to analyze the cleavage efficiency of the individual chains within the heterotrimer. We used 2D gels with the first dimension analyzed in non-reducing conditions to separate cleavage intermediates and the second dimension analyzed in reducing conditions to separate the CP products from the individual α1 or α2 chains (Fig. 2B). This revealed that there was indeed a marked deficit of α1(I) in the first cleavage intermediate, leading to an α1:α2 ratio of 0.4 ± 0.1 instead of the ratio of 2 which would be expected if the two chains were cleaved with the same efficacy. This result shows that BMP-1 preferentially cleaves the α2(I) chain and that the cleavage sequence in procollagen I heterotrimer most often starts with the processing of this chain to create the first and most difficult break in the trimer (Fig. 2C). In the presence of PCPE-1, the α1:α2 ratio in the first cleavage intermediate increases to 0.7 ± 0.1 (Fig. 2B), suggesting that PCPE-1 can help to partially overcome the cleavage bias but that its action is not sufficient to fully restore the 2:1 stoichiometry.

**Figure 2:**
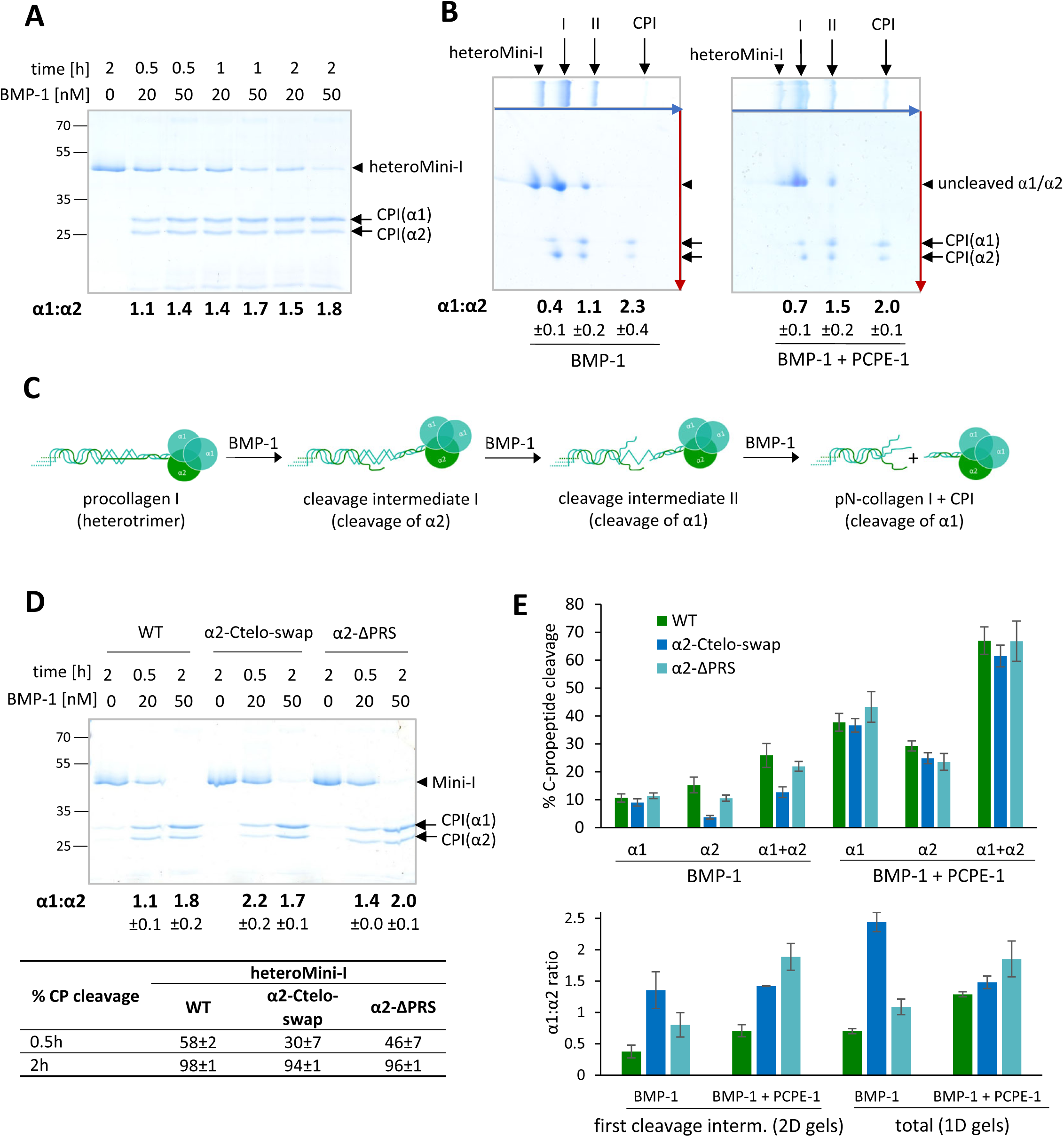
Comparison of the cleavage by BMP-1 of the α1(I) and α2(I) chains within heteroMini-I. **(A)** SDS-PAGE analysis (12 % acrylamide gel; reducing conditions) of the relative amounts of CPI(α1) and CPI(α2) generated at two BMP-1 concentrations (20 or 50 nM) and three time points (0.5, 1 and 2 h). The α1:α2 ratio was calculated from the densitometric quantification of the CP band intensities (one representative experiment; n = 3 independent experiments). **(B)** 2D SDS-PAGE analysis of the CPs of individual α chains in each of the cleavage intermediates. HeteroMini-I (900 nM) was incubated with 10 nM BMP-1 for 5 min (with PCPE-1, 900 nM) or 60 min (without PCPE-1). The first dimension (blue arrow) was in non-reducing conditions and the second dimension (red arrow) was in reducing conditions. I and II refer to successive cleavage intermediates derived from heteroMini-I, as also described in (C). The α1:α2 ratio is the mean ± SD of n = 3 independent experiments resulting from the densitometric quantification of CP dot intensities. **(C)** Proposed cleavage sequence for procollagen I processing by BMP-1. **(D)** SDS-PAGE analysis (12 % acrylamide gel; reducing conditions) comparing the relative amounts of CPI(α1) and CPI(α2) in heteroMini-I WT, α2-Ctelo-swap and ΔPRS after 0.5 h (20 nM BMP-1) or 2 h (50 nM BMP-1). The percentage of cleavage and the α1:α2 ratio represent the mean ± SD of n = 4 independent experiments. **(E)** Graphs showing the % of C-propeptide cleavage and the α1:α2 ratios for WT and mutant heteroMini-I when they are incubated with BMP-1, in the presence or absence of PCPE-1. Cleavage assays were run in conditions of low basal BMP-1 activity (4 nM BMP-1 for 0.5 h with 450 nM heteroMini-I and PCPE-1 for 1D gels; conditions like in (B) for 2D gels) and analyzed in 1D or 2D SDS-PAGE, as illustrated in Fig. S4A-B. Quantification of band intensities yielded the parameters for the first cleavage intermediate (from 2D gels) or for the average of processing intermediates and final products respectively (from 1D gels). Means ± SD of n = 3 (WT) or 4 (α2-Ctelo-swap and ΔPRS) independent experiments.

We then wanted to identify the specific features of the α2(I) chain which makes it more susceptible to BMP-1 proteolysis. Considering the canonical cleavage sites, both chains are cleaved between alanine and aspartate residues and the cleavage sequence is relatively well-conserved between the two α chains except for positions P3’-P5’ which correspond to an insertion of three residues (PRS) in the α2(I) chain (Fig. 1E). This can be important as prolines were previously reported to occur more frequently than any other amino-acid in the P3’ position of BMP-1 cleavage sites (38). In addition, the C-telopeptide sequence located on the N-terminal side of the cleavage site is 11-residue shorter in α2(I) than in α1(I) (Fig. 1E). This could lead to the α2(I) chain presenting a more extended conformation (Fig. 1A) and explain why it is more susceptible to proteolysis. To test the effect of these differences, we deleted the PRS sequence in α2(I) (α2-ΔPRS mutant; Fig. S1 and S3) and swapped the telopeptide of α2(I) with the telopeptide of α1(I) (α2-Ctelo-swap mutant; Fig. S1 and S3) in heteroMini-I. With BMP-1 alone, in conditions of both intermediate (Fig. 2D) or low (Fig. 2E; Fig. S4A) basal activity, the most striking effect was observed for the α2-Ctelo-swap mutant which led to a marked reduction in the level of cleavage of the α2 chain. This led to a decrease in overall cleavage of heteroMini-I by BMP-1 by around 50 % (CPIα1 + CPIα2; Fig. 2E) and allowed a very good correction of the preferential cleavage (the α1:α2 ratio was 2.2 at early time points in 1D gels and 1.4 in the first cleavage intermediate in 2D gels; Fig. 2D-E and Fig. S4A-B). The heteroMini-I containing the α2-ΔPRS mutation achieved a more modest correction of the cleavage bias with an α1:α2 ratio of 1.4 in 1D gels and of 0.8 in 2D gels. Together, this suggests that the C-telopeptide sequence is the most important factor to explain the efficient cleavage of the α2 chain, most likely through the tension/stretching of the cleavage site induced by the shorter length of the α2 C-telopeptide. Interestingly, when both mutants were cleaved in the presence of PCPE-1, the total activity was the same as for WT heteroMini-I (Fig. 2E) and the enhancement of α2 processing (EF(α2)) for α2-Ctelo-swap was significantly increased compared to WT and α2-ΔPRS (more than 3-fold; Fig. S4C). In contrast, the enhancement of α1 processing is comparable for all three proteins. This maintained the correction of the preferential cleavage in the first intermediate around 1.4 for α2-Ctelo-swap in the presence of PCPE-1 (Fig. 2E, Fig. S4B), similarly to the condition with BMP-1 alone. Here however, a more pronounced effect was observed for α2-ΔPRS with a full correction of the cleavage bias in 2D gels (α1:α2 ratio of 1.9; Fig. 2E). This suggests that the presence of the PRS sequence in the native α2 chain impedes the efficient enhancement by PCPE-1 or, alternatively, mimics the effect of PCPE-1 and abolishes the need for enhancement. Despite the significant differences observed with both mutant mini-procollagens in the presence of PCPE-1, their binding profiles to PCPE-1 were minimally altered compared to WT (Fig. S4D), again indicating that there is no direct correlation between binding affinity and efficacy of enhancement.

### Structure of heteroMini-I in complex with PCPE-1

As previous attempts to solve the crystal structure of recombinant heterotrimeric CPI were unsuccessful (2), we employed cryo-electron microscopy (cryo-EM) to investigate the 3D structure of heteroMini-I. All our attempts to obtain a high-quality model of heteroMini-I alone failed, likely due to the high flexibility of the triple helical domain, the relatively small molecular size, and the preferred particle orientation within the ice. To increase the effective size and limit conformational heterogeneity, we introduced PCPE-1 as a binding partner. This approach also presented significant challenges, as no clear density corresponding to PCPE-1 was initially observed under standard vitrification and imaging conditions. Only when an additional nickel-affinity enrichment step was performed, starting from the complex isolated by SEC, was PCPE-1 detected, appearing in 2D class averages as a diffuse density adjacent to the flower-like shape of the C-propeptide (Fig. S5A).

The C-propeptide of heteroMini-I adopts the characteristic flower-like architecture previously described for CPI and CPIII homotrimers (2, 13), comprising three petal-like lobes, a base, and a stalk (Fig. 3A). Each interface between the three chains is stabilized by calcium ions at the level of the base region. The overall binding mode of the PCPE-1 CUB domains also closely resembles that observed in the CPIII/PCPE-1 complex (31), with CUB1 predominantly interacting with residues in one α1 chain and CUB2 interacting with residues in the counterclockwise adjacent α chain (here α2). These interactions are mediated by key lysine residues from the individual collagen chains, K35 (α1) and K21 (α2) (numbering for CP chains starting at the canonical BMP-1 cleavage sites, as in previous reports (2, 13)), which form electrostatic contacts with calcium-coordinating residues within the respective CUB domains (Fig. 3A; insets). Additional stabilizing interactions are provided by residues Glu60, Tyr67, Asp68, Asp109, Gly111, and Thr112 in CUB1 of PCPE-1, and Glu183, Tyr190, Asp191, Asp233, Ser235, and Val236 in CUB2. The previously reported critical role of Phe90 in CUB1 (31, 41) was also confirmed; its side chain inserts between the two collagen α chains, forming a hydrophobic core that stabilizes the PCPE-1/CPI interface (Fig. 3A; lower inset).

**Figure 3:**
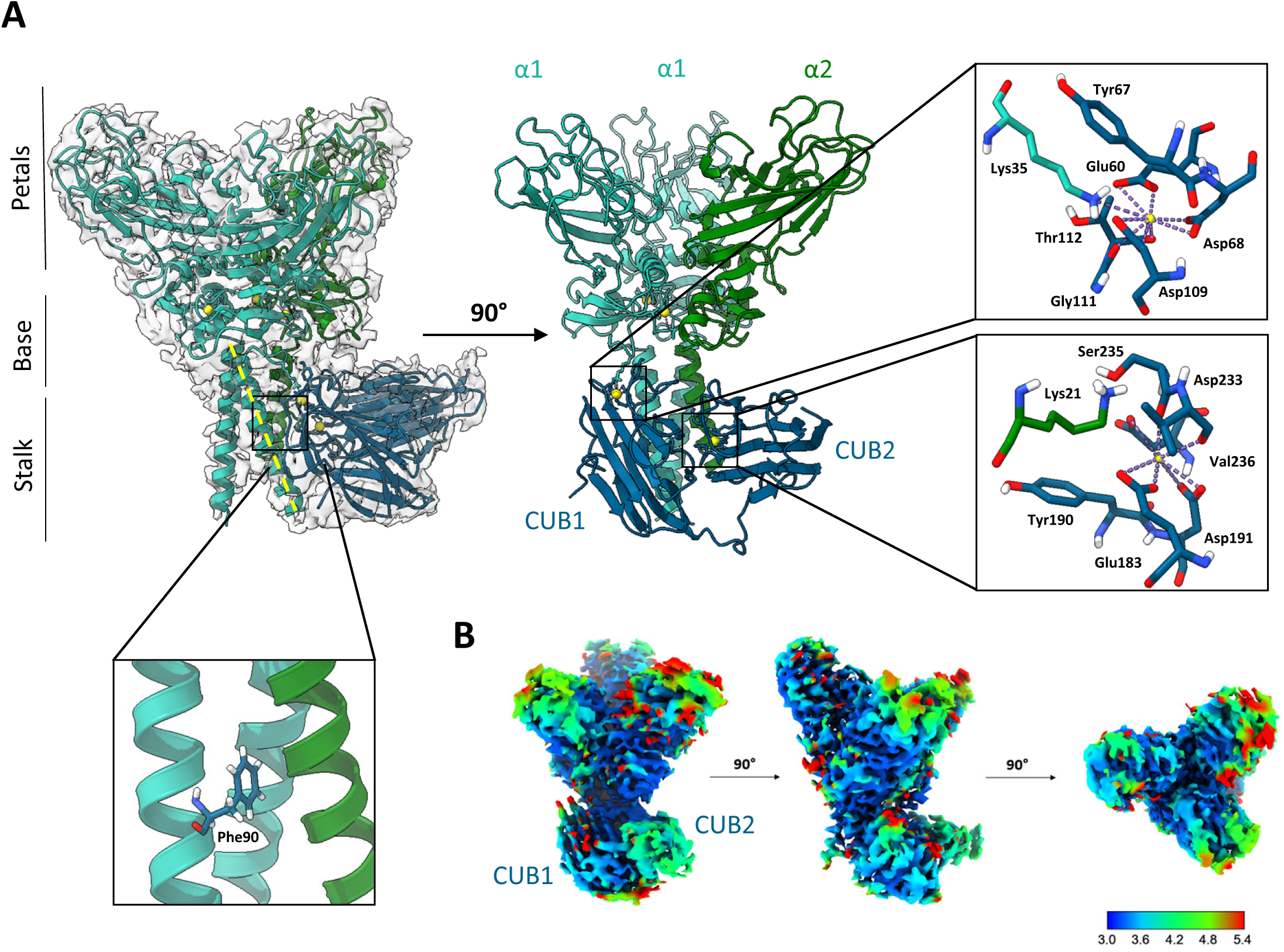
Cryo-EM structure of heteroMini-I with PCPE-1. **(A)** Cartoon representation and map-to-model fit (upper left panel) of the cryo-EM structure of heteroMini-I (α1 chains, cyan; α2 chain, green) complexed with PCPE-1 (marine blue). Close-up views show residues at the interaction interfaces: the upper right panel displays the α1 chain bound to the CUB1 domain of PCPE-1 through K35, and the lower right panel shows the α2 chain bound to the CUB2 domain through K21. Calcium ions are shown in yellow. The lower left panel highlights the positioning of PCPE-1 Phe90 between the coiled-coil helices of α1 and α2 chains. The yellow dotted line shows the absence of bending in the α1 helical stalk interacting with CUB1. **(B)** Local resolution map of the cryo-EM reconstruction, with blue indicating the highest and red the lowest local resolution. The map was generated using CryoSPARC v4.7.1.

Due to its assymetric nature, the structure of heteroMini-I in complex with PCPE-1 is expected to represent a mixture of the three possible binding modes of PCPE-1 to heterotrimeric CPI, which is reflected in relatively low CCmask and CCvolume values (0.57 and 0.56, respectively). However, the cryo-EM density was best modeled by the configuration where CUB1 interacts with one α1 chain and CUB2 with the α2 chain, as described above. Chain assignment was primarily based on differences in coiled-coil length, in particular the presence of Pro11 in the α2 chain, which acts as a helix breaker and shortens the ordered helical region, as clearly visible in the cryo-EM density. Independent validation of chain assignment was provided by binding affinity measurements of PCPE-1 with various heteroMini-I mutants (see below), supporting that the observed binding mode corresponds to the most stable and populated complex species.

Local resolution analysis revealed that the least well-resolved regions correspond to the most C-terminal regions of the α chains, composed mainly of flexible loops (“petals”), as well as to the CUB2 domain of PCPE-1 (Fig. 3B). A similar pattern of disorder in CUB2 was previously observed in the X-ray structure of CPIII in complex with PCPE-1 CUB domains, where nearly half of the CUB2 domain could not be modeled due to flexibility (31). Consistently, neither the triple-helical region and C-telopeptide of heteroMini-I nor the long linker and NTR domain of PCPE-1 were visible in the final reconstruction.

### Role of individual binding configurations in the heteroMini-I/PCPE-1 complex

In order to validate the existence of three binding modes for PCPE-1 on heterotrimeric procollagen I, despite the overrepresentation of the configuration described in Fig. 3A, we next prepared mutants of heteroMini-I which were designed to direct PCPE-1 to one specific interaction site. This was achieved through the specific removal of one or two of the lysines which are known to be critical for complex formation (Fig. 4A; Fig. S1). The corresponding mutants were obtained with the same protocol as described above for heteroMini-I and tested in binding and activity assays. First, we confirmed the critical role of K35 and K38 in α1(I) and α2(I) respectively with a heterotrimer combining 2 x α1-K35Q and 1 x α2-K38Q chains. This triple mutant had a very low binding affinity for PCPE-1 by SPR and poor enhancing activity in cleavage assays (Fig. 4B-C; Fig. S6A, C). Yet, the cleavage efficiency with BMP-1 alone is exactly the same as for the WT in conditions of both low and intermediate basal cleavage (Fig. 4C; Fig. S6A-B). This confirms that the mutations specifically affect the activity of PCPE-1, as is also the case for the other mutants discussed below.

**Figure 4:**
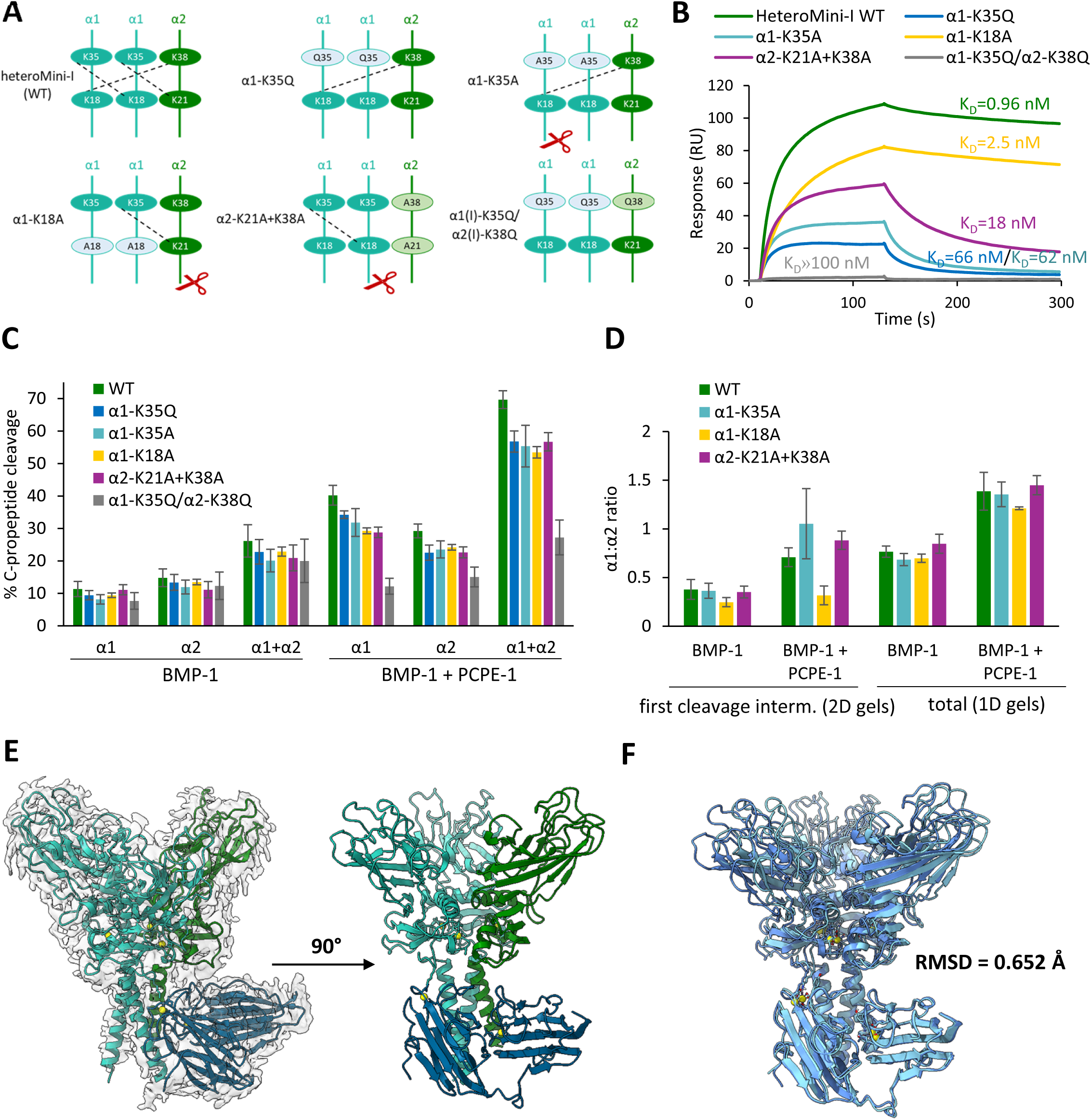
Evaluation of the individual contributions of the three binding modes to heteroMini-I/PCPE-1 complex formation and to enhancing activity. **(A)** Design of mutants to evaluate the role of each binding site in heteroMini-I. Dotted lines represent the expected position of PCPE-1 CUB domains (CUB1 engaged through K35/K38 and CUB2 through K18/K21). The red scissors indicate the chain that is cleaved first by BMP-1 upon PCPE-1 binding. **(B)** Comparison of the SPR interaction profiles of heteroMini-I and its mutants with PCPE-1. 50 nM of WT and mutants were injected over immobilized PCPE-1 (140 RU). **(C)** Graph showing the % of C-propeptide cleavage for WT and heteroMini-I binding mutants when they are incubated with BMP-1, in the presence or absence of PCPE-1. Cleavage assays were run in conditions of low basal BMP-1 activity (4 nM BMP-1 for 0.5 h with 450 nM heteroMini-I and PCPE-1), analyzed in 1D gels, as illustrated in Fig. S6A and quantified. Means ± SD of n = 5 (WT), n = 4 (α1-K35Q, α2-K21A+K38A and α1-K35Q/α2-K38Q), n = 3 (α1-K18A) and n = 6 (α1-K35A) independent experiments. **(D)** α1:α2 ratios for WT and heteroMini-I binding mutants in conditions of low basal BMP-1 activity like in (B) for 1D gels and in conditions of Fig. 2B for 2D gels. Quantification of band intensities yielded the ratios for the first cleavage intermediate (from 2D gels, Fig. S6D-I) or for the average of processing intermediates and final products respectively (from 1D gels, Fig. S6A). Means ± SD of n = 3 (WT, α1-K35A and α1-K18A) or n = 4 (α2-K21A+K38A) independent experiments for 2D gels or as described in (B) for 1D gels. **(E)** Cartoon representation and map-to-model fit (left panel) of the cryo-EM structure of the α1-K18A heteroMini-I (α1 chains, cyan; α2 chain, green) complexed with PCPE-1 (marine blue). **(F)** Superposition of WT (light blue) and α1-K18A (dark blue) structures of heteroMini-I complexed with PCPE-1, showing an RMSD of 0.652 Å.

If mutation of K35 in α1 (α1-K35Q or α1-K35A) is combined with native α2, it is expected that CUB1 will bind to heteroMini-I through α2 only and CUB2 through α1 only, as depicted in Fig. 4A. Strikingly, for these mutants, we observed a lower stability of the complex than for the WT (Fig. 4B; Fig. S7) and yet they displayed a similar level of enhancing activity for both α chains (Fig. S6C). Still, in 2D gels (Fig. 4D; Fig. S6D-E), α1-K35A leads to a slightly better correction of the preferential cleavage of α2 than the WT, in the presence of PCPE-1, suggesting that it may favor the cleavage of the α1 chain. For this mutant, the α1 chain for which enhanced cleavage is observed can be either the α1 chain binding CUB2 or the second α1 chain not involved in PCPE-1 binding (Fig. 4A). In both cases, this rules out the main model proposed by Pulido *et al.* (31) suggesting that the chain interacting with CUB1 (here α2; Fig. 4A) is the chain that is cleaved first by BMP-1 in the presence of PCPE-1. In order to determine which of the two other chains (binding CUB2 or free) is the target of BMP-1 upon PCPE-1 binding, we analyzed the mutants directing PCPE-1 to the two other binding sites (α1-K18A and α2-K21A/K38A; Fig. 4A and Fig. S6F-I). While binding occurs through two α1 chains in α2-K21A+K38A, this mutant had a similar behavior to α1-K35Q and α1-K35A in terms of binding and cleavage parameters, suggesting that the chain cleaved preferentially in the first cleavage intermediate is also α1 for this mutant. In contrast, α1-K18A has a much higher affinity for PCPE-1 (close to WT; Fig. 4B; Fig. S7) but is completely unable to correct the cleavage bias (Fig. 4C). This clearly indicates that this mutant, which directs the binding of PCPE-1 to α1 via CUB1 and to α2 via CUB2, further promotes the cleavage of α2 when PCPE-1 is bound. Together, the results establish that the chain that is cleaved first upon PCPE-1 binding is the chain engaged through CUB2 (Fig. 4A, red scissors) and not through CUB1 as previously proposed (31). Also, we can conclude that the three binding configurations of the heteroMini-I/PCPE-1 complex are possible and active. However, the configuration observed with the α1-K18A mutation has a higher affinity which explains why it was more prominent in the cryo-EM data collected for the WT. This also suggests that complex formation *in vivo* may not result in equal proportions of the three binding modes and may vary with the conditions, especially the relative concentrations of procollagen I and PCPE-1.

To improve map quality and increase CC values, an additional structure of the α1-K18A mutant in complex with PCPE-1 was determined (Fig. S8). This reconstruction yielded significantly improved CCmask and CCvolume values (0.82 and 0.80, respectively) and confirmed that a single binding mode can be experimentally isolated (Fig. 4E). Superposition of the wild-type (WT) and α1-K18A structures showed near-identical conformations, with an RMSD of 0.652 Å, indicating that the overall architecture of the complex is conserved (Fig. 4F).

### Comparison of trimer interfaces in heterotrimeric and homotrimeric C-propeptides

To compare heterotrimeric CPI with homotrimeric CPI, the CPI coordinates were extracted from the heteroMini-I/PCPE-1 structure (PDB ID: 9SHT) and superimposed with the homotrimeric CPI structure from a previous study (PDB ID: 5K31) (2). The comparison revealed a highly similar overall fold, with an RMSD of 1.789 Å across 4745 aligned atoms (Fig. 5A), indicating that the global architecture of the C-propeptide trimer is conserved between homo- and hetero-trimeric procollagen I.

**Figure 5:**
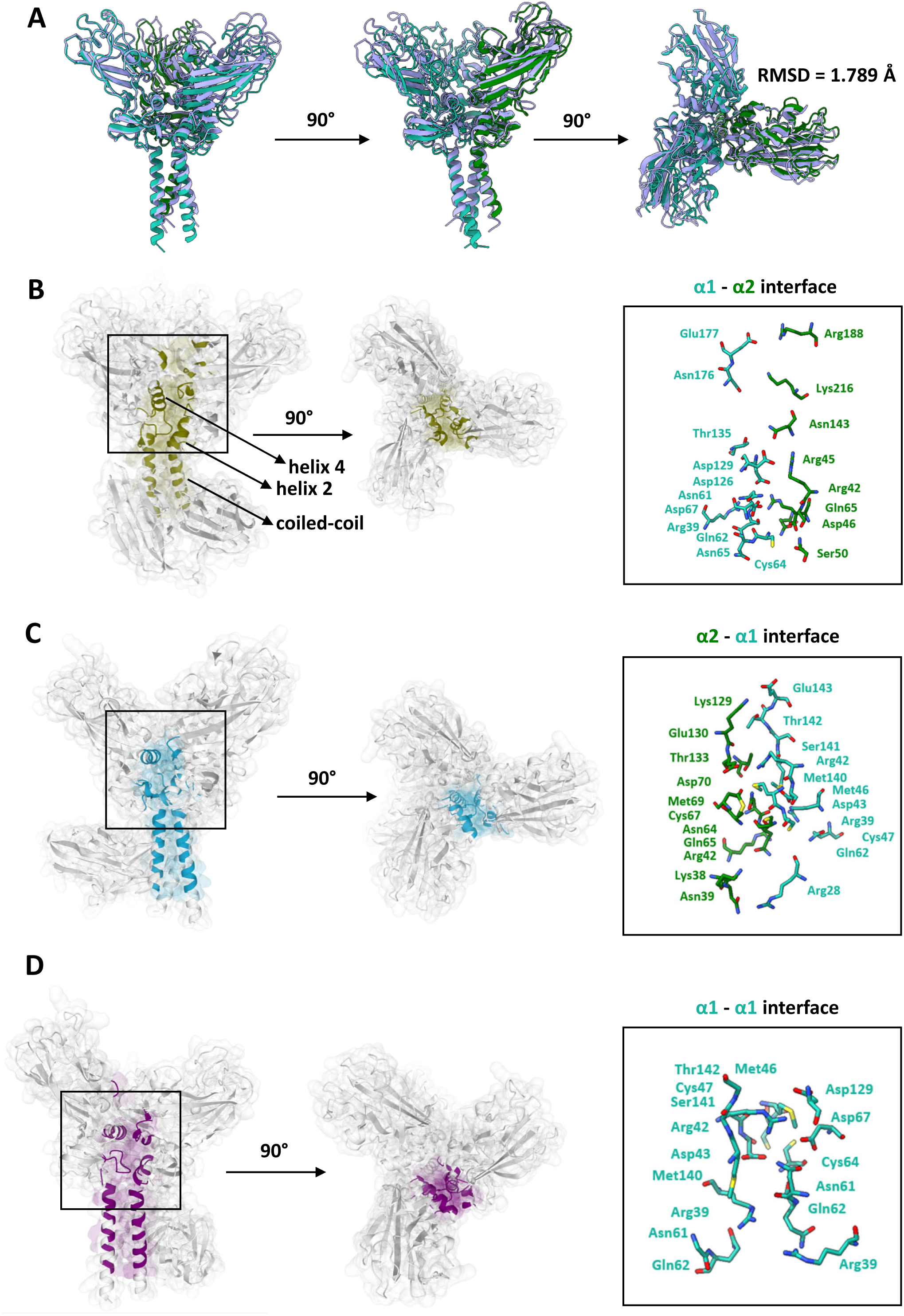
Structural comparison of heterotrimeric and homotrimeric collagen I C-propeptides and analysis of interchain interfaces in the heterotrimer. **(A)** Superposition of the heterotrimeric CPI structure from this study (α1, cyan; α2, green), extracted from the WT heteroMini-I/PCPE-1 complex (PDB ID: 9SHT), with the homotrimeric CPI structure (PDB ID: 5K31; grey-blue). Structural alignment was performed using the MatchMaker tool in UCSF ChimeraX and optimized for maximal agreement within the coiled-coil region of the C-propeptides. (B) Schematic representation of the interchain interface between the α1 chain interacting with the CUB1 domain and the α2 chain interacting with the CUB2 domain of PCPE-1, generated using the MAPIYA Contact Maps server (left panel), together with a close-up view highlighting charged residues at the interface. (C) Schematic representation of the interchain interface between the α2 chain and the α1 chain not interacting with PCPE-1, generated using the MAPIYA Contact Maps server (left panel), together with a close-up view highlighting charged residues at the interface. (D) Schematic representation of the interface between the two α1 chains, generated using the MAPIYA Contact Maps server (left panel), together with a close-up view highlighting charged residues at the interface. All interface analyses were performed using the α1-K18A heteroMini-I/PCPE-1 model. Close-up views exclude residues involved in interactions within the coiled-coil regions of the individual α chains.

The structures described in this study also enable detailed analysis of the interfaces between procollagen α chains in the heterotrimer. Due to the improved map quality and clearer definition of the α1-K18A heteroMini-I/PCPE-1 complex, all subsequent interchain interface analyses were performed using this structure. Global analysis of interchain contacts using the MAPIYA Contact Maps server (42) demonstrated that trimerization is primarily mediated by interactions involving the coiled-coil regions, helix 2, helix 4, and the loop preceding helix 4 of the procollagen α chains (Fig. 5B–D, left panels). The analyzed interface areas ranged from 826 to 941 Å², with the α1-α2 and the α2-α1 interfaces having higher areas than the α1-α1 interface (Table S2). The latter exhibited the smallest buried surface area (826 Å²), closely matching the interfaces observed in the homotrimeric CPI structure (PDB ID: 5K31), which range from 821 to 843 Å². However, the calculated Gibbs free energy changes (ΔG) do not follow the same order with the α1-α2 interface now having the lowest value, indicating that interface area does not directly correlate with the overall strength of interactions. Importantly, in the α1-K18A structure, disulfide bridges were not modeled because no corresponding density was observed in the cryo-EM maps. Nevertheless, their presence is strongly supported by SDS-PAGE analysis, where addition of DTT strongly reduced the apparent molecular weight of the trimer (Fig. S3). The absence of visible disulfide bonds in the reconstruction may result from radiation-induced bond breakage during electron exposure or from insufficient local map quality to confidently resolve these features. Notably, homoMini-I contains three intermolecular disulfide bridges, one at each interchain interface, while heteroMini-I contains only two. The α1-α2 interface is indeed devoid of interchain disulfide bond (14) and this may contribute to the mild destabilization of the heterotrimer, as suggested by the manual inclusion of disulfide bonds in ΔG calculations which now increases the difference with the other interfaces up to 20-30 % (Table S2). Another important aspect is that the contribution of calcium ions to the stabilization of the interface is not taken into account in the ΔG calculations made with PDBePISA which only considers apo forms. However, calcium ions are known to play a critical role to stabilize the C-propeptide chain interfaces in the first steps of the trimerization process (14) and a recent molecular dynamics study suggested that it is only in the presence of calcium that the interchain interactions become significantly stronger in the heterotrimer compared to the homotrimer (43). Taken together, our results show that the presence of the α2 chain does not substantially favor the thermodynamic stability of the procollagen I trimer at the inter-residue level. However, additional stabilizing (calcium ions) and destabilizing (absence of one intermolecular disulfide bridge) factors must be considered to explain the predominance of heterotrimers over homotrimers *in vivo*.

Interchain interactions at the C-propeptide level are known to initiate the process of procollagen trimerization and are primarily mediated by salt bridges and hydrogen bonds between neighboring chains. Within the α1-α1 interface, the previously reported importance of residues Arg42 and Asp129 in stabilizing the interface was confirmed (2) as both residues participate in hydrogen-bond and salt-bridge formation. Additional contributions from Asn61, Asp67, Arg39, and Asp43 were also identified (Fig. 5D and Table S2). In the α1-α2 interface, involved in PCPE-1 binding in our structures, we could not confirm the previously proposed interaction between Asp129 of α1 and Arg45/Lys247 of α2. This interface also exhibited the lowest number of hydrogen bonds and salt bridges among all analyzed interfaces. Notably, the unpaired Cys64 residue of the α2 chain formed interactions with Asp46 and Ser50 of α1, while Asn61 interacted with Arg42 (Fig. 5B and Table S2). At the α2-α1 interface, our structure confirmed the interaction between Lys129 and Glu143, as well as the involvement of Arg42 from the α1 chain (2). In contrast, no direct interaction between this residue and Glu130 of the α2 chain (2), was observed although its side chain is positioned near the interface. However, additional stabilizing contributions were identified for Thr133 and residues located at the N-terminus of the so-called “chain recognition sequence” (CRS; (13)) region of the α2 chain (Fig. 5C and Table S2).

### The “chain unraveling” mechanism previously proposed for PCPE-1 does not apply to heteroMini-I

Comparison with the CPIII/PCPE-1 complex structure (PDB ID: 6FZW) (31) likewise demonstrated conservation of the overall structural organization despite local differences, yielding an RMSD of 4.942 Å across 4538 aligned atoms (Fig. S9A). Similarly, superposition of heteroMini-I with the apo CPIII structure (PDB ID: 4AK3) (30) produced an RMSD of 1.902 Å over 3857 aligned atoms (Fig. S9B). Together, these analyses demonstrate that the overall architecture of CPI and CPIII is highly conserved and that PCPE-1 binding does not induce major conformational rearrangements within the trimeric assembly.

Intriguingly, a notable difference between the heteroMini-I/PCPE-1 structure presented here and the previously reported CPIII/PCPE-1 structure is the absence of any detectable bending or distortion within the α1 chain engaged by the CUB1 domain of PCPE-1. In the CPIII/PCPE-1 complex, formation of a salt bridge involving Asp5 of the collagen chain and residues within CUB1 was proposed to promote local chain unraveling and to facilitate access of the protease to the procollagen cleavage site (31). Asp 5 is not conserved in any of the CPI chains and both heteroMini-I WT and α1-K18A structures show a well-ordered and linear coiled-coil region within the CUB1-engaged α1 chain, with no evidence of local distortion or unwinding (Fig. 3A; inset in Fig. S10A). These observations suggest that the previously proposed PCPE-1 enhancement mechanism involving chain unraveling around the P5’ position of the cleavage site does not apply to heterotrimeric procollagen I and may not be the main driver of PCPE-1 activity.

## Discussion

In the present study, we describe the successful production of heterotrimeric and homotrimeric mini-procollagens I. To our knowledge, this is the first description of the expression and purification of a recombinant heterotrimeric (mini-)procollagen I allowing the precise control of the correct 2:1 stoichiometry. As we observed around 12 % of homotrimer co-expressed with the heterotrimer, it is likely that some homotrimer contamination was also present in previous studies describing the recombinant production of heterotrimeric collagen I (18–20). Indeed, the pepsin extraction that is commonly used to purify collagen I does not allow the use of tags for affinity purification and for the control of chain composition. Thus, our system is unique in that it allows the precise control of chain assembly and composition. Our analysis of the thermal denaturation of heteroMini-I and homoMini-I also confirmed the slightly higher thermal stability of the homotrimer, as previously described for the corresponding C-propeptides (2) and mature collagens (11), and the calcium dependence of the two proteins for proper folding (14).

Our constructs also proved extremely versatile, allowing for a broad range of functional and structural characterizations. First, from the cryo-EM data collected for the complex between heteroMini-I and PCPE-1, we could derive the first high-resolution structure for heterotrimeric CPI and compare it to the previously described structure of homotrimeric CPI (2). The two proteins have a similar flower-like fold with three petals and an almost perfect conservation of the C-propeptide architecture. Although the area of the interfaces involving the α2 chain were slightly larger than the α1-α1 interfaces found in homo- or hetero-trimers, the α1-α2 interface is overall less thermodynamically stable due to the lack of one interchain disulfide bond. Therefore, there is no obvious benefit in terms of trimer stabilization to integrate the α2 chain and the large prevalence of heterotrimers observed *in vivo* must result from another type of regulation. Indeed, a recent molecular dynamics study by Johnson *et al.* suggested that preferential association with the α2 chain requires prior calcium binding and that the calcium-bound heterotrimer has stronger interchain contacts than the calcium-bound homotrimer (34).

Another interesting application of the mini-procollagens described here is the determination of cleavage sites for C-proteinases and the possibility to make semi-quantitative kinetic experiments. In this context, we were able to confirm the previously reported cleavage sites for BMP-1 but also to reveal one additional cleavage in the α1 chain of procollagen I, occurring some residues downstream of the canonical site, which had never been described before. A similar secondary cleavage site for BMP-1 was also recently reported in a mini-procollagen II substrate (44) with *in vivo* evidence that this second cleavage occurs in the native procollagen II molecule (45). Cleavage site identification could be further extended in the future to the other PCPs such as mTLL1 (46), meprins (21), ADAMTS-2 or ADAMTS-14 (28, 47).

Mechanistic studies are now also possible with heterotrimeric procollagen I, especially through site-directed mutagenesis, to characterize the molecular determinants of its interactions with proteolytic enzymes or various potential partners such as PCPE-1. Here, we have characterized seven new mutants in both α1 or α2 chains and derived extremely valuable information regarding the mechanism of procollagen I processing. First, we have solved a long-standing controversy regarding the order of cleavages in heterotrimeric procollagen I (48, 49) as we could unambiguously establish that the α2 chain is cleaved first in the majority of the heterotrimers. The presence of this chain actually provides a striking advantage in terms of cleavage efficiency compared to the homotrimer. As this chain is processed much faster by BMP-1, the first break in the trimer is easier to make and the subsequent cleavages of α1 chains in the partially disrupted trimer are then facilitated. This explains the striking two-fold increase in BMP-1 activity observed when heteroMini-I was compared to homoMini-I and suggests that the higher resistance to proteolysis is a general property of (pro)collagen homotrimers. Indeed, it was previously reported that the triple helix of homotrimers is also more difficult to cleave by human collagenases (12).

Two important features of the α2 chain were identified as plausible molecular determinants explaining its efficient processing. First, the short length of the α2 C-telopeptide is expected to promote kinks in the looser α1 C-telopeptide and to stretch the N-terminal side of the cleavage site in the α2 chain, thereby potentially facilitating the attack of this chain by the protease. Secondly, the insertion of the PRS sequence on the prime (C-terminal) side of the α2 cleavage site, possibly in collaboration with the two other prolines located some residues further, causes a premature interruption of the α-helical coiled-coil organization of this chain whereas, in α1 chains, the cryo-EM structures clearly show that the coiled coil domain extends to the cleavage site. We can thus anticipate that the disruption of the well-structured coiled-coil in α2 facilitates the cleavage by the protease. Proline-rich motifs are also frequently associated with protein-protein interaction modules and conformational regulation (50, 51), raising the possibility that the PRS sequence participates in the selective recruitment or positioning of the protease. Interestingly, the PRS sequence also affects the enhancing activity of PCPE-1, making the α2 chain “resistant” to stimulation. Although the molecular basis for this finding remains unclear (the PRS sequence is unfortunately not visible in the cryo-EM structures), it is tempting to speculate that the role of PCPE-1 is to reproduce the stretching/distortion effects naturally occurring within the α2 chain.

Our attempts to obtain structural information on other two-partner (mini-procollagen/BMP-1) or three-partner (mini-procollagen/BMP-1/PCPE-1) complexes have been unsuccessful so far. However, the results obtained in this study allow us to propose a new model to explain how PCPE-1 can enhance the cleavage of fibrillar procollagens by BMP-1. Based on the X-ray structure of the complex between CPIII and the CUB domains of PCPE-1, we had previously proposed that PCPE-1 could function by locally unwinding the coiled-coil domain of CPIII to isolate the distorted chain and facilitate its cleavage by BMP-1 (31). Unexpectedly, this distortion is not observed in the model derived from the cryo-EM data of the heteroMini-I/PCPE-1 complex. As mentioned above, the coiled-coil region of the α1 chain interacting with CUB1 is well-defined down to position P2’ of the canonical cleavage site and does not show any sign of bending even beyond the point where unraveling is supposed to occur. This observation clearly indicates that the mechanism previously proposed to explain the stimulation of procollagen processing by PCPE-1 is unlikely to represent a universal mode of PCPE-1 activity, at least in the context of heterotrimeric procollagen I.

Another conclusion from the present work is that the chain that is cleaved first by BMP-1, when PCPE-1 is bound to procollagen I, is the chain engaged through the CUB2 domain of PCPE-1. Intriguingly, CUB2 is also the CUB domain with the lowest resolution in the cryo-EM structures suggesting that it is less tightly bound to the procollagen than CUB1. The less stable interaction is also supported by previous SPR results suggesting a lower affinity of CUB2, compared to CUB1, for mini-procollagen III (40). Nevertheless, CUB2 is able to direct the protease to the chain where it is bound, suggesting that the dynamic nature of this interaction can be an advantage when it comes to accommodating the protease in the three-partner complex. Our new hypothesis is thus that the CUB2 domain of PCPE-1 could be involved in a specific interaction with BMP-1, most probably through its non-catalytic domains. Along this line, it is interesting to note that the CUB2 domain of the cognate PCPE-2 protein was recently shown to establish strong contacts with the CUB2 domain of BMP-1 (52), although this tight interaction leads to a different outcome (inhibition) in the BMP-1/PCPE-2 complex (29, 53). The hypothesis that a CUB2/CUB2 interaction could also be involved in the mechanism of stimulation by PCPE-1 is supported by the fact that PCPE-1 cannot stimulate the cleavage of procollagen I by BMP-1 when the CUB2 domain of the protease is missing (Lipinski *et al.* in preparation), clearly indicating that this domain is playing a major role in the enhancing activity of PCPE-1. However, this interaction is expected to occur only in the presence of the procollagen substrate as we have previously demonstrated that the binding of BMP-1 to PCPE-1 alone is weak (29, 54) and that the interaction of BMP-1 with mini-procollagen III is stabilized in the presence of PCPE-1 (30). Together, our results point to a model where CUB1(PCPE-1) anchors PCPE-1 to one chain of the procollagen substrate while CUB2(PCPE-1), that is loosely bound to an adjacent procollagen chain, attracts CUB2(BMP-1) in order to (i) induce a conformational change in this specific chain or (ii) anchor the protease in the optimal position for efficient cleavage or (iii) both. In all cases, these mechanisms would facilitate procollagen cleavage by the catalytic domain of BMP-1. The nature of the potential conformational change that could be triggered through this CUB2/CUB2 interaction remains to be elucidated but, based on the α2 mutants described here, the possible stretching of the cleavage site, through distant anchoring points (involving the catalytic and CUB2 domains of BMP-1), appears as a plausible option.

Importantly, the remaining collagen chain not directly engaged by PCPE-1 does not appear to undergo efficient activation despite being more solvent-exposed. This observation suggests that productive cleavage requires precise spatial organization rather than simple accessibility of the cleavage site. Thus, cooperative interactions between procollagen I, PCPE-1, and BMP-1 define a preferred geometric and kinetic regime that governs productive catalysis. Unexpectedly, we have also observed that PCPE-1 activity is less sensitive to the nature of procollagen chains and trimeric assemblies than to the level of basal cleavage by BMP-1. The proteins and mutants which were less efficiently cleaved by BMP-1 (*e.g.* homoMini-I, α2-Ctelo-Swap) were also extremely well stimulated by PCPE-1, regardless of the chain, binding affinity and trimeric context, suggesting that PCPE-1’s primary function is to assist the formation of the first break in the trimer while subsequent cleavages are minimally affected. This would explain why the effect of PCPE-1 becomes less visible in conditions of high basal BMP-1 activity when most substrate molecules have already passed the step of the challenging first cleavage event. This also helps to rationalize the observation that the enhancement factor tends to decrease over the course of the reaction and requires low basal cleavage for appropriate evaluation. Collectively, our results show that PCPE-1 acts as a kinetic and structural organizer that modulates both the spatial arrangement and the sequence of substrate engagement.

In conclusion, together with the previously described mini-procollagens II and III (23, 29, 44), the novel mini-procollagens I form a valuable toolkit to characterize the process of collagen assembly, both at the level of initial trimer formation (through CP-driven chain recognition) and at the final level of fibril formation (through PCP-driven proteolytic processing). We found that they were very useful in mechanistic and structural studies, thanks to their high purity and versatility in mutagenesis studies, and expect many more future applications.

## Materials and methods

### Production of mini-procollagens I

The DNA sequences encoding the two chains of mini-procollagen I with supplementary GPP repeats and tags but without signal peptide, as described in Fig. S1, were synthesized by Genewiz using native sequences with mutations of N-glycosylation sites. Integration into the pHLsec vector was also done by Genewiz using the AgeI and XhoI cloning sites, in frame with the signal peptide already included in the vector (55). Insert sequences were verified by Sanger sequencing (Eurofins) before plasmid amplification in *E. coli* XL1-Blue bacteria. Plasmids were then purified using the NucleoBond PC 10000 EF Giga kit for endotoxin-free plasmid DNA from Macherey-Nagel.

In parallel, HEK293-F cells were grown in FreeStyle 293 expression medium (Gibco) (31) using sterile flasks (Corning or BD Biosciences) placed on an orbital shaker platform rotating at 125 rpm at 37 °C with 8 % CO_2_ (Eppendorf). On the day of transfection, cells were centrifuged and resuspended at a cell density of 1.10^6^ cells/mL in FreeStyle 293 expression medium. For transfection, the ratio of total DNA (in a 2:1 mass ratio for the α1 and α2 chains of the heterotrimer) to cells was 1 µg DNA for 10^6^ cells and the ratio of total DNA to PEI Max transfection agent (Polyethylenimine 25 K linear or MAX 40 K linear, Polysciences) was 1 µg DNA for 3 µg transfection agent. Plasmid DNA and PEI Max transfection agent were first diluted separately in 1/20 of the total culture volume of Opti-MEM medium (Gibco), filtered on 0.2 µm filters (Millipore) for the transfection agent and kept at room temperature for 5 min before mixing. The mixture was further incubated for 15 min before addition to the cells. 2-phospho-L-ascorbic acid trisodium salt (100 µM final concentration) was also added to the culture. Finally, 1/5 of the culture volume of fresh medium was added to the cells 24 hours after transfection.

Five days after transfection, the suspension was centrifuged at 1,000 g for 10 min, the cells were discarded and *N*-ethyl-maleimide (Merck, 2 mM) and Pefabloc (Roth, 0.25 mM) were added to the culture medium before centrifugation for 10 min at 10,000 g, 4° C. The pH of the supernatant was adjusted to 8. If the α2(I) chain was present, a biotin-blocking solution (1 mL in 500 mL of supernatant; IBA lifesciences) was added and incubated with the medium for 15 min at 4 °C under gentle stirring. The cell supernatants were frozen at -80 °C until purification.

### Purification of mini-procollagens I

The purification of heteroMini-I was achieved in two steps using the Twinstrep tag on the α2 chain and size exclusion chromatography. After equilibration of the Strep-tactin XT Superflow resin (IBA lifesciences, 10 mL per liter of cell supernatant) in 0.1 M Tris-HCl pH 8, 0.3 M NaCl, the cell supernatant was loaded at 0.5 mL/min and 4 °C. The column was washed with 5 volumes of equilibration buffer and eluted with 6 volumes of the same buffer supplemented with 50 mM D-biotin (IBA lifesciences). The fractions containing heteroMini-I were pooled and concentrated by ultrafiltration (Vivaspin 20, 30 kDa, Sartorius) before injection in a HiLoad 10/300 Superdex S200 Increase gel filtration column (Cytiva) equilibrated in 20 mM HEPES pH 7.4, 0.15 M NaCl, 5 mM CaCl_2_. Useful fractions were concentrated around 1 mg/mL and stored at -80 °C until further use.

The purification of homoMini-I was achieved in three steps using Affi-Gel Blue Gel, Butyl Sepharose, and size exclusion chromatography. In the first step, the Affi-Gel Blue Gel (Bio-Rad) column was equilibrated in 20 mM HEPES, 150 mM NaCl, 2 mM CaCl_2_, pH 7.4, followed by application of the clarified supernatant at 0.5 mL/min and 4 °C. Elution was performed using the same buffer supplemented with 2 M NaCl or 50 % ethylene glycol. The eluted fractions containing homoMini-I were subjected to the second step employing hydrophobic interaction chromatography on Butyl Sepharose (Bio-Rad). The sample was adjusted to approximately 1.2 M (NH₄)₂SO₄ and loaded into the resin equilibrated in 1 M (NH₄)₂SO₄, 50 mM Tris, pH 8. Elution was achieved by a stepwise decrease in salt concentration, and fractions containing homoMini-I were pooled and concentrated by ultrafiltration (Vivaspin 20, 30 kDa, Sartorius). In the final step, protein was purified by gel filtration on a HiLoad 10/300 Superdex S200 Increase gel filtration column (Cytiva) equilibrated in 20 mM HEPES, 150 mM NaCl, 5 mM CaCl_2_, pH 7.4. Fractions corresponding to the main peak were pooled, concentrated to approximately 1 mg/mL and stored at -80 °C until further use.

### Other proteins and mutants

Human BMP-1 (with a Flag-tag at the C-terminus) and human PCPE-1 (with 8His-tag at the C-terminus for cleavage assays and cryo-EM or without tag for other experiments) were obtained as described before (23, 41, 56).

Mutations in α1(I) or α2(I) were introduced by site-directed mutagenesis using primers (Microsynth) carrying the desired nucleotide substitution. Due to difficulties in amplifying the pHL-sec vector directly by PCR, the procedure was divided into two stages. In the first stage, only the gene insert encoding the respective chain was amplified. Two overlapping PCR fragments were generated – one spanning from the 5’ end to the mutation site and the other from the 3’ end to the mutation site, so that both products shared a region containing the introduced mutation. In each of these reactions, one primer carried the point mutation, while the opposite primer included a restriction site compatible with the cloning strategy. The overlapping fragments were then used as templates in a subsequent PCR to obtain full-length double-stranded insert. The complete insert and the pHL-sec vector were digested with AgeI and XhoI restriction enzymes (New England Biolabs). The digested products were excised from agarose gels and ligated using Quick Ligase (New England Biolabs). The resulting plasmids were transformed into *E. coli* DH5α cells, and plasmid DNA was isolated from selected colonies (Macherey-Nagel). The constructs were verified by sequencing (Microsynth), and plasmids containing the expected mutations were used in subsequent experiments.

All the heteroMini-I mutants were purified using the same protocol as for the WT heteroMini-I.

### Pepsin digestion

One µg heteroMini-I or homoMini-I in 5 µL of 20 mM HEPES, 150 mM NaCl, 5 mM CaCl_2_, pH 7.4 was acidified with 1 µL of 0.5 M acetic acid to reach a final pH of 2.5. Pepsin (P6887, Merck) was dissolved at 1 mg/mL in 0.5 M acetic acid and 0.5 µL was added to the previous solution as indicated. The mixture was left for 2 h or overnight at RT in static conditions. For optimal detection of the triple helical domain, proteins were labeled with the Cy5 dye, using the QuickStain protein labeling kit (Amersham), according to the manufacturer’s instructions. Samples were boiled for 5 min in electrophoresis sample buffer and analyzed by SDS-PAGE. Visualization was with a Typhoon gel imager (Amersham) at 600 V.

### Western blotting

After SDS-PAGE, proteins were transferred to polyvinylidene difluoride (PVDF) membranes by liquid transfer in 10 mM *N*-cyclohexyl-3-aminopropanesulfonic acid (pH 11) with 10 % ethanol (2 h, 280 mA) and blocked for 2 hours in 10 % (w/v) skim milk in PBS. Primary antibodies against PCPE-1 (1:10,000) and C-propeptide α1(I) (LF-41; 1:1,000) were home-made rabbit polyclonal antibodies, prepared as described in (23) and (57) respectively. The Streptactin coupled to alkaline phosphatase was from IBA Lifesciences. The antibodies were diluted in 5 % skim milk, 0.05 % Tween 20 in PBS and applied on membranes overnight at 4 °C. After three washes in 0.05 % Tween 20 in PBS, horseradish peroxidase (HRP)-coupled anti-rabbit secondary antibodies (Cell Signaling Technology, 1:10,000) were added for 1 h at room temperature. Signal was detected, after extensive washing of the membrane with 0.05 % Tween 20 in PBS, using the enhanced chemiluminescence technique with the ECL Select kit (Cytiva) and the FX Fusion camera (Vilber Lourmat) or the colorimetric detection with the AP Conjugate Substrate kit (Biorad).

### Cleavage assays

HeteroMini-I or homoMini-I were incubated with BMP-1, in the presence of absence of PCPE-1 (with an 8His-tag at the C-terminus (41)), at 37 °C under agitation (350 rpm) for the time indicated in figure and table legends. The cleavage buffer was 50 mM HEPES pH 7.4, 150 mM NaCl, 5 mM CaCl_2_, 0.02 % octyl β-D-glucopyranoside. Standard reaction mixtures for 1D gel analyses contained 450 nM Mini-I, 450 nM PCPE-1, and BMP-1 at the concentration indicated in the figure legends. The reactions were started by the addition of the protease and stopped with 25 mM EDTA and/or SDS-PAGE sample buffer, followed by boiling for 8 min at 95 °C. They were analyzed by SDS-PAGE on 4-12 % acrylamide gradient gels (Thermo Scientific) or home-made gels with acrylamide concentrations adapted to the reaction conditions, and visualized with InstantBlue Coomassie Protein Stain (Abcam) or by ATOMS. Alternatively, analysis could also be performed using two-dimensional electrophoresis, as described below. As the N-terminal cleavage products are not stained very clearly, the percentage of BMP-1–dependent cleavage was calculated using the following formula: 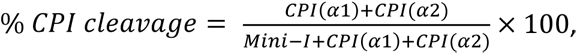 where the values refer to the intensity of corresponding bands (Mini-I or CPI) on SDS-PAGE. % cleavages for CPI(α1) or CPI(α2) were calculated using only the relevant CPI value in the numerator. EF (Enhancement Factor) with PCPE-1 (total) was calculated using the formula: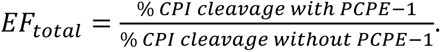 EF for specific chains (α_1_ or α ) was calculated using the formula: 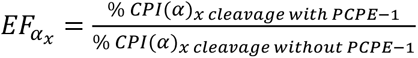 where (α)_x_ refers to the relevant α chain.

### 2D electrophoresis

Two-dimensional SDS-PAGE was performed following the protocol described in (58) with minor modifications. Prior to electrophoresis, cleavage assays were performed with or without PCPE-1. Reactions without PCPE-1 were incubated for 60 minutes, and those with PCPE-1 for 5 min, both at 37 °C and 350 rpm. Each reaction (30 µL total volume) contained 900 nM Mini-I, 900 nM PCPE-1, and 10 nM BMP-1 in cleavage buffer (50 mM HEPES, 150 mM NaCl, 5 mM CaCl_2_, 0.02 % octyl β-D-glucopyranoside, pH 7.4). Reactions were stopped by adding SDS loading buffer without DTT and boiled for 8 min at 95°C. Samples were loaded into first-dimension 8 % polyacrylamide gels (1 mm thick). After electrophoresis, the gel strips were incubated for 15 minutes at room temperature in 0.1 M DTT diluted in SDS-PAGE running buffer, rinsed, and then incubated for another 15 minutes in the dark with 0.1 M iodoacetamide in the same buffer. The second-dimension gels (15 %, 1.5 mm thick) were cast using ¾ of the usual volume of stacking gel to accommodate the first-dimension strips. Each strip was positioned horizontally (higher molecular weight to the left) on top of the second-dimension gel and overlaid with 0.5 % agarose in SDS-PAGE buffer containing 0.5 g/L bromophenol blue. A piece of Whatman paper containing the protein ladder was placed on the left side of each gel. Gels were run under standard SDS-PAGE conditions (not exceeding 200 mV), stained with fresh InstantBlue Coomassie Protein Stain (Abcam), and destained with water.

### NanoDSF

Thermal denaturation was analyzed by NanoDSF using a Prometheus NT.48 instrument (Nanotemper). HeteroMini-I and homoMini-I were diluted to a final concentration of 150 µg/mL in 20 mM HEPES pH 7.4, 150 mM NaCl, 5 mM CaCl_2_ (Ca^2+^ condition) or 20 mM HEPES pH 7.4, 150 mM NaCl, 20 mM EDTA (EDTA condition). Ten microliters of proteins were loaded into a capillary (PR-C002, Nanotemper) and thermal denaturation was recorded using a 1 °C/min ramp from 20 to 95 °C, with fluorescence detection at 330 nm and 350 nm (excitation at 280 nm). Data were recorded using the PR.ThermoControl v2.3.1 software and analyzed with the PR.StabilityAnalysis v1.1 software (Nanotemper). Melting temperatures were determined as the inflection points of the curves representing the ratio of fluorescence intensity at the two wavelengths (350nm/330nm) as a function of temperature.

### Surface Plasmon Resonance

Surface plasmon resonance experiments were run at 25 °C with a Biacore T200 equipment (Cytiva) equipped with the Biacore T200 control software (v3.2.1). Human PCPE-1 (no tag (23)) was diluted in 10 mM HEPES pH 7.4 and covalently immobilized on Series S sensor chips CM5 (Cytiva) using the reagents included in the Amine coupling kit (Cytiva). SPR signals were recorded simultaneously on a control channel where a mock immobilization procedure was performed (no protein ligand) and double referencing was applied to all injections. HeteroMini-I, homoMini-I and heteroMini-I mutants were injected at 50 μL/min after dilution in running buffer (10 mM HEPES pH 7.4, 0.15 M NaCl, 5 mM CaCl_2_ and 0.05 % P20) for 120 s. Regeneration was achieved with 2 M guanidinium chloride. Best fits of the data were computed with the Biacore T200 evaluation software (v3.2.1). In Fig. S7, a “weighted average KD” was also calculated according to the following formula:

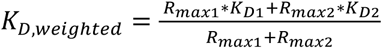

### ATOMS

Five micrograms of partially cleaved and uncleaved mini-procollagen samples, prepared as above, were processed for TMT-ATOMS, as previously described (36, 37) with minor modifications. Briefly, samples were denatured in 2.5 M guanidinium chloride and 1.8 M HEPES pH 8.0 at 65 °C for 15 min, reduced with 10 mM Tris (2-carboxyethyl) phosphine (TCEP) at 65 °C for 45 min, and alkylated with 25 mM iodoacetamide at room temperature in the dark for 30 min. After TMT labeling in DMSO (TMT label:protein mass ratio = 26), the mini-procollagen samples (alone, with BMP-1, with BMP-1 + PCPE-1) were mixed and concentrated using the SP3 (single-pot, solid-phase-enhanced sample-preparation) protocol (59). Proteins and SP3 beads (GE healthcare) were mixed in a 1:10 weight ratio in 80 % ethanol for 5 minutes at RT. Then, supernatant was discarded and beads were washed three times on a magnetic rack with 80 % ethanol before being resuspended in 100 µL of 180 mM HEPES pH 8. The supernatant was digested overnight at 37 °C with sequencing-grade trypsin or chymotrypsin (Promega; 1:100 protease:protein weight ratio). To remove SP3 beads, the samples were placed on a magnetic rack for 5 minutes, and supernatants were centrifuged at 14,000 g for 10 min. After desalting on a C18 column (Pierce) and drying under vacuum, the digested samples were dissolved in 0.1 % formic acid and analyzed by LC-MS/MS using a Q-Exactive HF mass spectrometer (Thermo Scientific; heteroMini-I experiments), equipped with a RSLC Ultimate 3000 nanoLC system (Thermo Scientific), or an Exploris 480 mass spectrometer (Thermo Scientific; homoMini-I experiments), equipped with a Vanquish NEO nano-LC system (Thermo Scientific). Data files were analyzed with Proteome Discover 2.4 or 2.5 using the SEQUEST HT search engine against a homemade database including the sequences of the mini-procollagens, BMP-1 and PCPE-1 sequences. Precursor mass tolerance was set at 10 ppm, fragment mass tolerance at 0.02 Da and up to 3 missed cleavages were allowed. Semi-tryptic digestion was selected. Oxidation (M, P), acetylation (Protein N-terminus), and TMT6Plex (N-term, K) were set as variable modifications and carbamidomethylation (C) as fixed modification. Peptides and proteins were filtered using the XCorr confidence threshold calculation based on the ion precursor charges and the MS/MS spectra. Quantification was performed with the Reporter Ions Quantifier node. The peak integration was set to the Most Confidence Centroid with 20 ppm Integration Mass Tolerance on the reporter ions. Protease/control ratios were calculated as the means of all the ratios obtained for N-terminally-labeled peptides defining cleavage sites (regardless of C-terminal length or modifications) or mature N-termini (after signal peptide removal). Ratios showing a fold change of at least two were conserved as defining cleavage sites in Table 2.

### Grid preparation for cryo-EM

To stabilize the heteroMini-I trimer and increase the effective particle size for cryo-EM analysis, PCPE-1 was used as a binding partner. Because standard purification conditions did not yield particles with clearly detectable PCPE-1 density, an additional nickel-affinity enrichment step was introduced prior to vitrification.

Purified heteroMini-I (both WT and α1-K18A mutant) and PCPE-1 carrying a C-terminal 8×His tag were mixed at a 1:2 molar ratio (200 µg Mini-I and 153 µg PCPE-1) and incubated on ice for 30 minutes. The mixture was applied to a Ni Spin Column (NEBExpress S1427S) equilibrated with binding buffer (20 mM HEPES, 150 mM NaCl, 2.5 mM CaCl_2_, pH 7.5). After washing with binding buffer containing 20 mM imidazole, the complex was eluted with elution buffer (same as binding buffer with 500 mM imidazole) and further purified by size-exclusion chromatography on a Superdex 200 Increase 10/300 column (Cytiva) equilibrated with 20 mM HEPES, 300 mM NaCl, 5mM CaCl_2_ and 0.02 % octyl β-D-glucopyranoside, pH 7.5. The main peak corresponding to the heteroMini-I/PCPE-1 complex (confirmed by SDS-PAGE) was concentrated to OD₂₈₀ = 0.2 using Vivaspin 20 concentrator (30 kDa MWCO, Sartorius).

Quantifoil R2/1 Cu 200 mesh grids, either without or with a 2 nm continuous carbon support layer, were glow-discharged prior to vitrification. Aliquots of 3 µL sample were applied to the grids and vitrified by plunge-freezing into liquid ethane using a Vitrobot operated at 4 °C and 100 % humidity, with a blot time of 5.5 s and blot force of −10. Grids were stored in liquid nitrogen until data collection.

### Cryo-EM Data acquisition and processing

Grid screening for the WT heteroMini-I/PCPE-1 complex was performed on a Glacios microscope (Thermo Scientific) at the Institute of Structural Biology (Grenoble, France) equipped with a Falcon4i direct electron detector. Screening data were collected at a nominal magnification of 150,000x, corresponding to a calibrated pixel size of 0.93 Å/pixel. Movies were collected with a total accumulated dose of ∼40 e⁻/Å² distributed over 45 frames and a defocus range from −0.8 to −2.2 µm.

Based on screening results, two grids were selected for high-resolution data collection: Quantifoil R2/1 Cu 200 mesh grids with and without a 2 nm continuous carbon support layer. High-resolution data collection was carried out at the CM02 beamline of the European Synchrotron Radiation Facility (Grenoble, France) using a Titan Krios G4 microscope operated at 300 kV and equipped with a Falcon4i detector and a Selectris X energy filter (10 eV slit width). Data were acquired at a nominal magnification of 165,000×, corresponding to a calibrated pixel size of 0.73 Å/pixel. Movies were fractionated into 42 frames with a total accumulated dose of 45 e⁻/Å² and collected within a defocus range of −0.8 to −2.4 µm. In total, 5,916 movies were collected from grids containing a continuous carbon support layer and 21,506 movies from unsupported grids.

For the α1-K18A heteroMini-I/PCPE-1 complex, high-resolution data collection was carried out on Quantifoil R2/1 Cu 200 mesh grid, with no preliminary screening, at the Oxford Particle Imaging Center using a Titan Krios G3i microscope operated at 300 kV and equipped with a Falcon4i detector and a Selectris X energy filter (10 eV slit width). Data were acquired at a nominal magnification of 165,000×, corresponding to a calibrated pixel size of 0.73 Å/pixel. In total, 10,109 movies were collected in lzw compressed tiff format, fractionated into 50 frames with a total accumulated dose of 50 e⁻/Å² and collected within a defocus range of −0.8 to −2.4 µm.

Data processing for the two heteroMini-I/PCPE-1 complexes was performed using CryoSPARC v4.7 (60).

Both datasets obtained with WT heteroMini-I were initially processed independently (Fig. S5B). Following Patch Motion Correction and Patch CTF Estimation, particles were initially picked using Blob Picker and subjected to template-free 2D classification. Resulting templates were subsequently used for template-based particle picking across both datasets. After additional rounds of 2D classification, particles displaying additional density corresponding to PCPE-1 were selected for ab initio reconstruction and heterogeneous refinement with three classes, followed by non-uniform refinement. The class exhibiting the best-defined secondary-structure features was subjected to reference-based motion correction and final refinement, yielding a reconstruction at 3.26 Å resolution according to the gold-standard FSC 0.143 criterion.

For the α1-K18A/PCPE-1 complex, following Patch Motion Correction and Patch CTF Estimation, particles were initially picked using Blob Picker/template picking and subjected to template-free 2D classification (Fig. S8B). After 2D classification, well-resolved classes were selected, and five ab initio models were generated to train a Topaz model and pick a new set of particles. A total of 770,000 particles belonging to the high-resolution class were selected and refined using NU-refinement with per-particle CTF refinement, leading to a 2.87 Å resolution map after reference-based motion correction. High-resolution heterogenous refinement with seven classes was performed leading to three high resolution maps featuring α1-K18A heteroMini-I/PCPE-1 complex. Particles belonging to the three classes were selected and subjected to high resolution ab-initio with 3 classes. One class containing 274,000 was refined using NU-refinement to 3.01 Å resolution. To minimize the intrinsic flexibility, the heteroMini-I petals and the PCPE-1/coiled coil of heteroMini-I were independently refined using local NU-refinement, leading to 2.91 Å and 2.97 Å maps respectively. The focus-refined maps were combined using PHENIX combine focused map (61) and used for model building after assessing that the quality of the combined map was identical to the focused maps and that the interface was identical to the consensus map.

Model building started from an initial model generated with AlphaFold 3 (62), consisting of two α1(I) chains and one α2(I) chain of procollagen I, full-length PCPE-1, and five calcium ions. The model was first fitted into the cryo-EM density map using rigid-body fitting in ChimeraX (63). Global restraints were generated at 6 Å, and all atoms were refined simultaneously in Coot (64). The resulting model was further refined in Phenix (61) using *readyset* to add hydrogen atoms and generate coordination for metal ions, followed by one round of real-space refinement.

Assignment of α1 and α2 chains within heteroMini-I was based on differences in the coiled-coil regions, including disruption of the α2 helix by Pro11. Due to conformational flexibility, the triple-helical region and C-telopeptide of heteroMini-I, as well as the linker and NTR domain of PCPE-1, were not resolved in the final reconstruction.

The final cryo-EM maps and the refined atomic model for the WT heteroMini-I/PCPE-1 complex were deposited in the Electron Microscopy Data Bank (EMDB) and Protein Data Bank (PDB) under accession codes EMDB ID: EMD-54910 and PDB ID: 9SHT, respectively. The final cryo-EM maps and the refined atomic model for the α1-K18A/PCPE-1 complex were deposited in EMDB and PDB under accession codes EMD-57262 (composite map), EMD-57259 (consensus map), EMD-57260 (focused map of the PCPE-1 region), EMD-57261 (focused map of the Mini-I petal regions), and PBD ID: 29OC, respectively.

## Acknowledgements

This work was inspired by the preliminary work made by David Hulmes (LBTI, UMR5305 Lyon). We thank Frédéric Delolme and Adeline Page from the Protein Science Facility of UMS 3444 (SFR Biosciences, Lyon) for performing mass spectrometry analyses, and Xavier Jaurand and Anton Sak from the Centre Technologique des Microstructures (University of Lyon, Villeurbanne) for the access to the cryo-EM grid preparation equipment. We also thank Céline Freton from MMSB (UMR 5086, Lyon) for providing access to the Prometheus NanoDSF apparatus and Pauline Juyoux for assistance with cryo-EM. We acknowledge the Oxford Particle Imaging Center (OPIC) for access to electron microscopy equipment. This work also used the EM facilities at the Grenoble Instruct-ERIC Center (ISBG; UMS 3518 CNRS CEA-UGA-EMBL) (65) with support from the French Infrastructure for Integrated Structural Biology (FRISBI; ANR-10-INSB-05-02) and GRAL, a project of the University Grenoble Alpes graduate school (Ecoles Universitaires de Recherche) CBH-EUR-GS (ANR-17-EURE-0003) within the Grenoble Partnership for Structural Biology. The IBS Electron Microscope facility is supported by the Auvergne Rhône-Alpes Region, the Fonds Feder, the Fondation pour la Recherche Médicale and GIS-IBiSA. Finally, we acknowledge the support from the CNRS/IN2P3 Computing Center (Lyon - France) for providing computing and data-processing resources needed for the cryo-EM analysis.

---

**Supplementary Figure S1:**
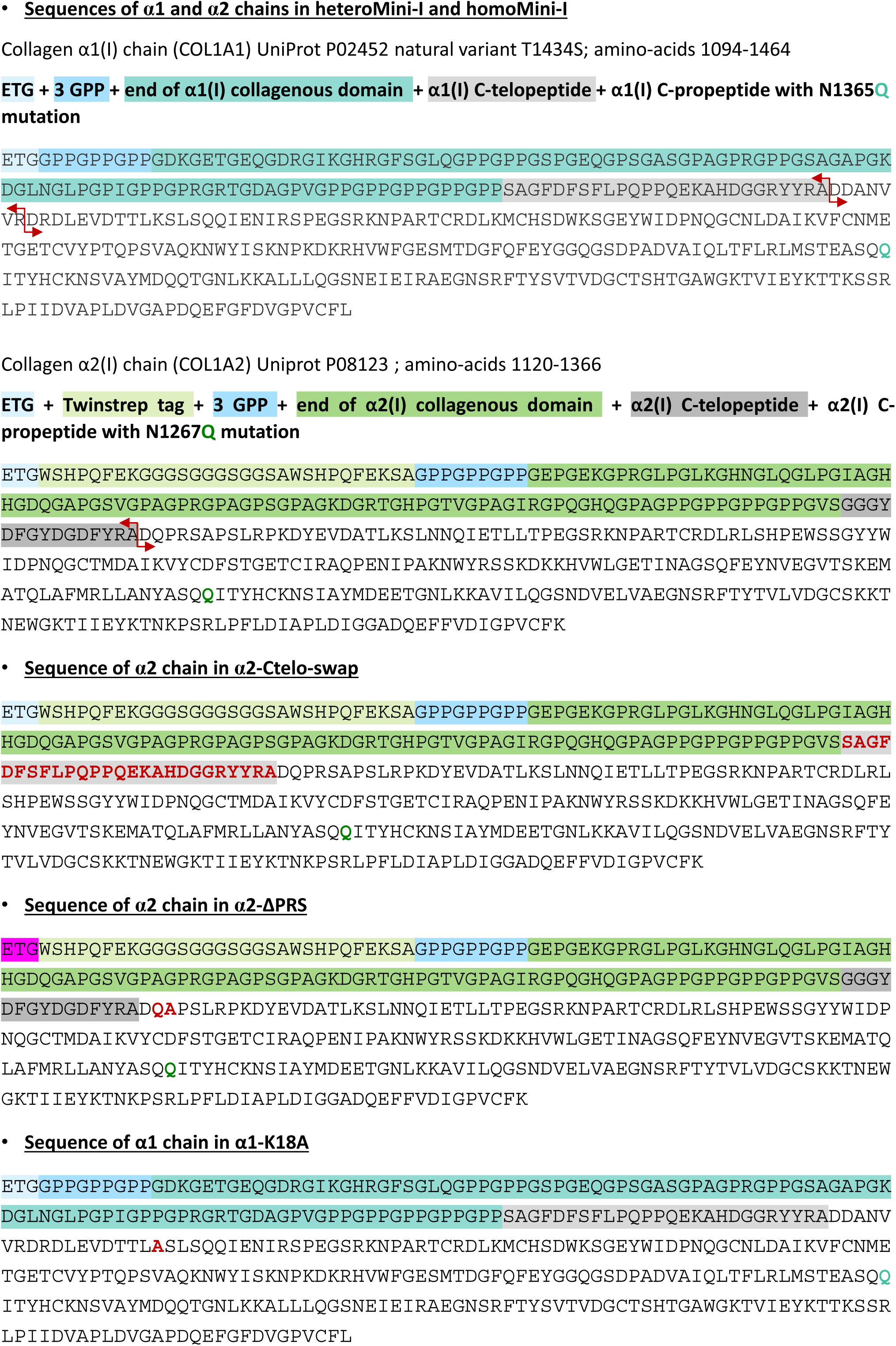
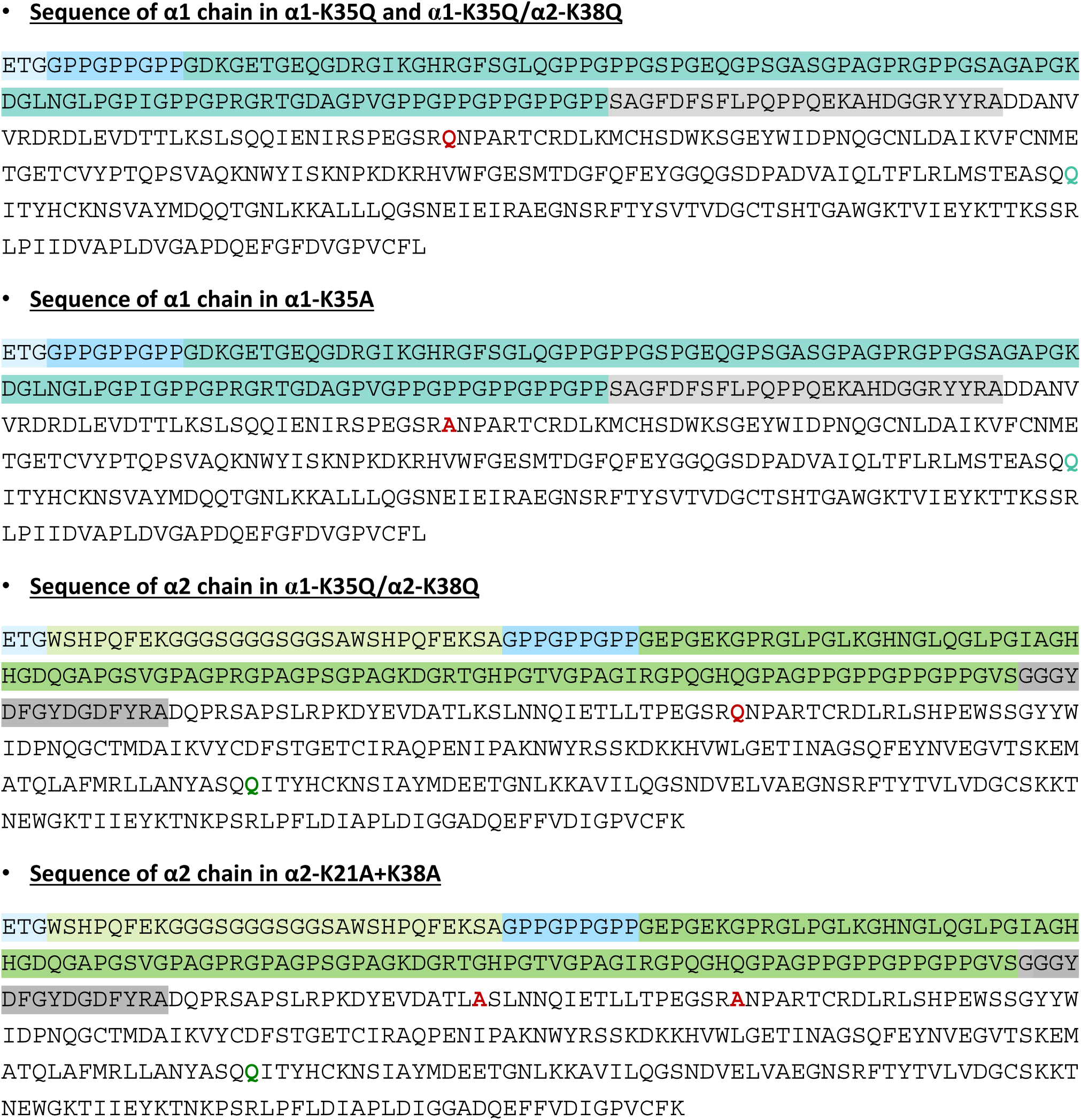
Mature protein sequences for the mini-procollagens (α1(I) and α2(I) chains) and their mutants. The different parts of the sequences are shown in different colors (color code as indicated above sequences); the cleavage sites for BMP-1, identified in this study, are indicated by red arrows; mutated residues are indicated in red color.

**Supplementary Figure S2:**
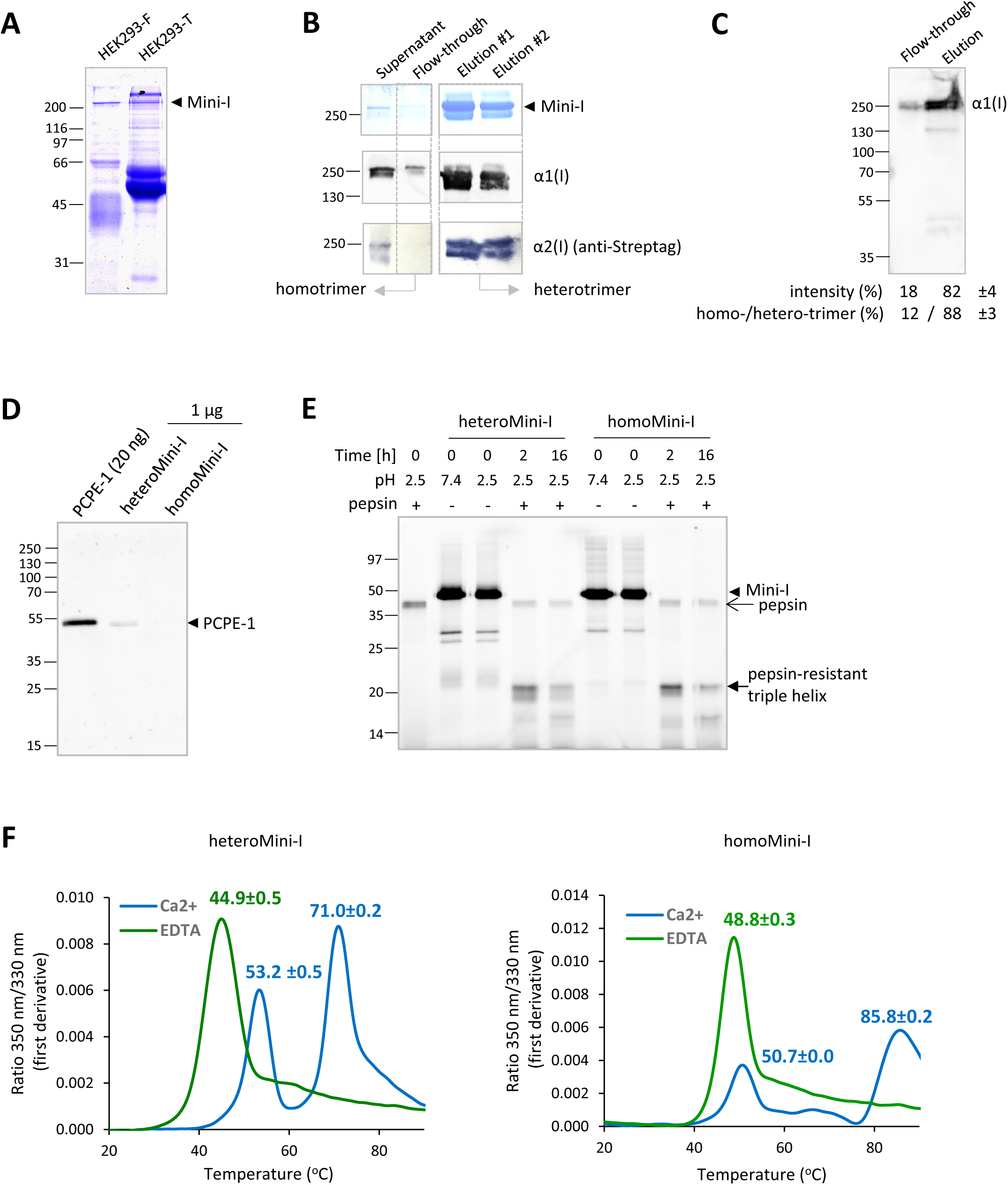
Production, purification and quality control of heteroMini-I and homoMini-I. **(A)** SDS-PAGE analysis of the conditioned medium of HEK293-T and HEK293-F after transfection with pHL-sec plasmids encoding the α1 and α2 chains of heteroMini-I (10 % acrylamide gel, non-reducing conditions, Coomassie Blue staining). **(B)** SDS-PAGE and western blot analysis (both in non-reducing conditions) of the composition of 293-F cell supernatant, collected 4 days after transfection of the plasmids encoding the α1(I) and α2(I) constructs, and flow-through and elution fractions of the Strep tactin XT Superflow column. The heteroMini-I trimers were detected after InstantBlue staining, the α1(I) chain with the LF41 antibody and the α2(I) chain with Streptactin-AP. **(C)** Immunodetection of the α1 chain in equivalent volumes of the flow-through and elution fractions with the LF41 anti-CPI(α1) antibody. Quantification of the relative proportions of homotrimers (in flow-through) and heterotrimers (in elution) based on signal intensity (Means ± range of two independent experiments). A correction was applied to account for the different numbers of α1 chains in hetero- and homo-trimers. **(D)** Immunoblot showing PCPE-1 detection in the final batches of heteroMini-I and homoMini-I after purification. One microgram of each mini-procollagen was loaded on a 10 % acrylamide gel (non-reducing conditions) and recombinant human PCPE-1 (20 ng) was used as positive control. **(E)** Pepsin digestion of heteroMini-I and homoMini-I was used to check the correct folding of the triple helix. The mini-procollagens (1 µg/sample) were incubated at pH 2.5, the optimal pH for pepsin, or at pH 7, as a control, in the absence or presence of 0.5 µg pepsin for 0, 2 or 16 h and the resulting products were analyzed by SDS-PAGE (15 % acrylamide; reducing conditions) after Cy5 labeling detection, as described in the Materials and Methods section. Control lanes with pepsin alone or mini-procollagens alone are also shown. **(F)** Analysis of the thermal stability of mini-procollagens in the presence and absence of calcium by NanoDSF. The fluorescence emitted at 350 nm and 330 nm was recorded during the thermal denaturation of the mini-procollagens and the first derivatives of the 350 nm / 330 nm ratios were calculated for heteroMini-I and homoMini-I. Curves are means of n = 7 (heteroMini-I with calcium or EDTA), n = 3 (homoMini-I with calcium) or n = 4 (homoMini-I with EDTA) independent experiments performed in duplicate. The corresponding melting temperatures (Tm; means ± SD) are indicated above the curves.

**Supplementary Figure S3:**
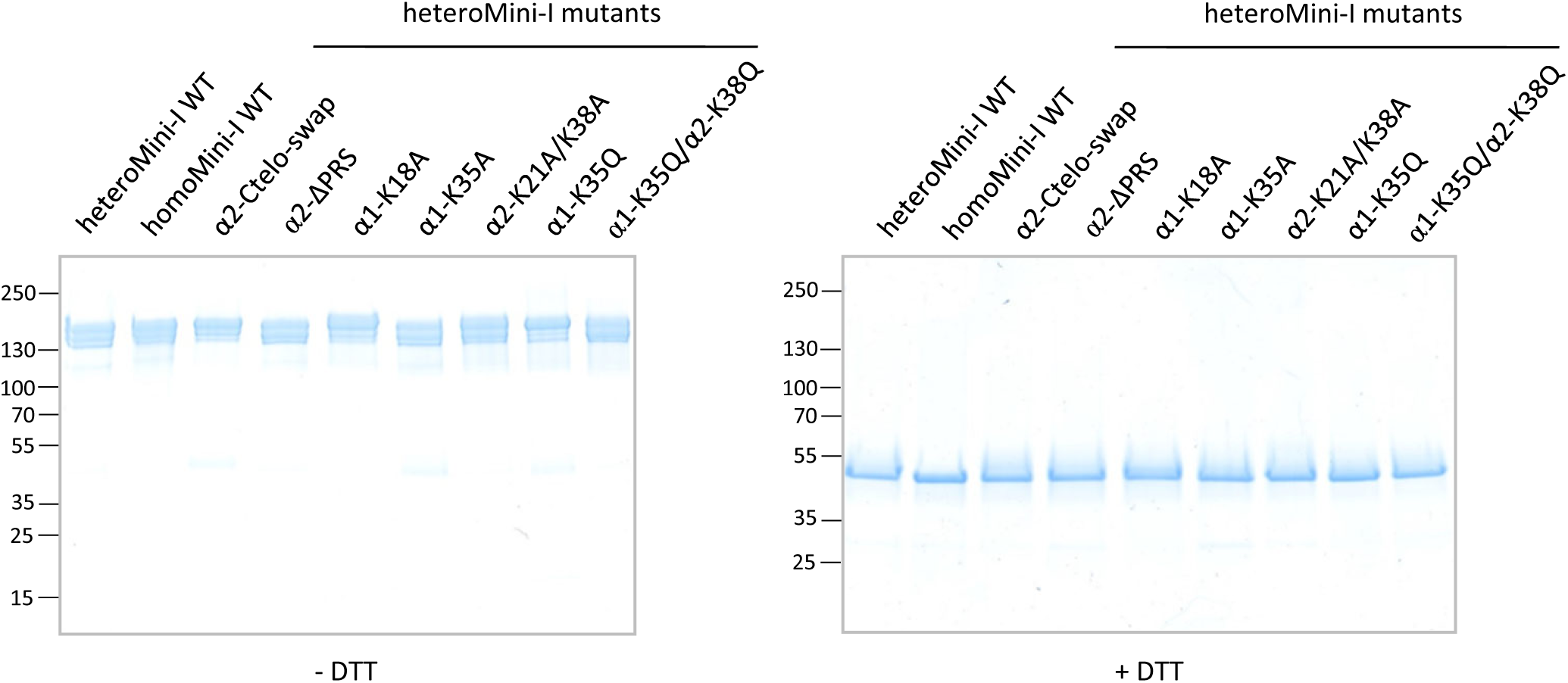
Purity of mutant heteroMini-I. SDS-PAGE analysis of the various mutant proteins produced in this study (4-20 % acrylamide; non-reducing - left and reducing conditions - right).

**Supplementary Figure S4:**
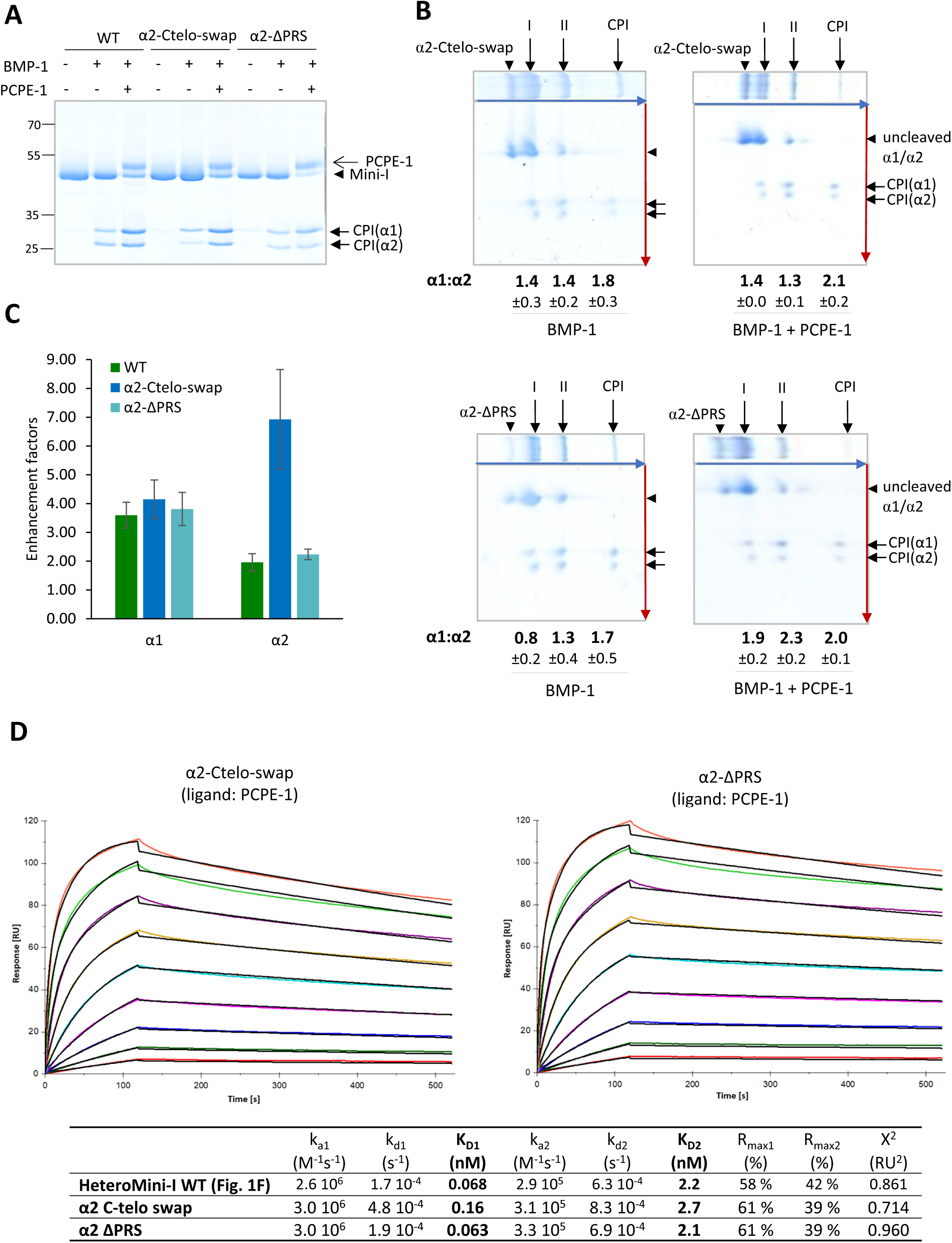
1D and 2D gels of cleavage assays and SPR analysis of interaction with PCPE-1 for α2-Ctelo-swap and α2-ΔPRS mutants. **(A)** 1D SDS-PAGE analysis (11 % acrylamide gel, reducing conditions) or **(B)** 2D SDS-PAGE analysis of the cleavage products of individual α chains in WT, α2-Ctelo-swap or ΔPRS when they are incubated with BMP-1 in the presence or absence of PCPE-1. The experimental conditions are the same as in Fig. 2E for (A) and Fig. 2B for (B). **(C)** Enhancement factors were calculated in conditions of low basal BMP-1 activity from 1D-gel densitometric analysis. **(D)** Interaction of α2-Ctelo-swap and α2-ΔPRS with immobilized PCPE-1 (140 RU). Increasing concentrations of the mutant proteins (0.39–100 nM; serial two-fold dilutions, including a duplicate injection at 6.25 nM) were injected. The best fits of the experimental sensograms obtained using the heterogeneous ligand model are shown (black curves), and the corresponding kinetic and dissociation constants are provided in the table below.

**Supplementary Figure S5:**
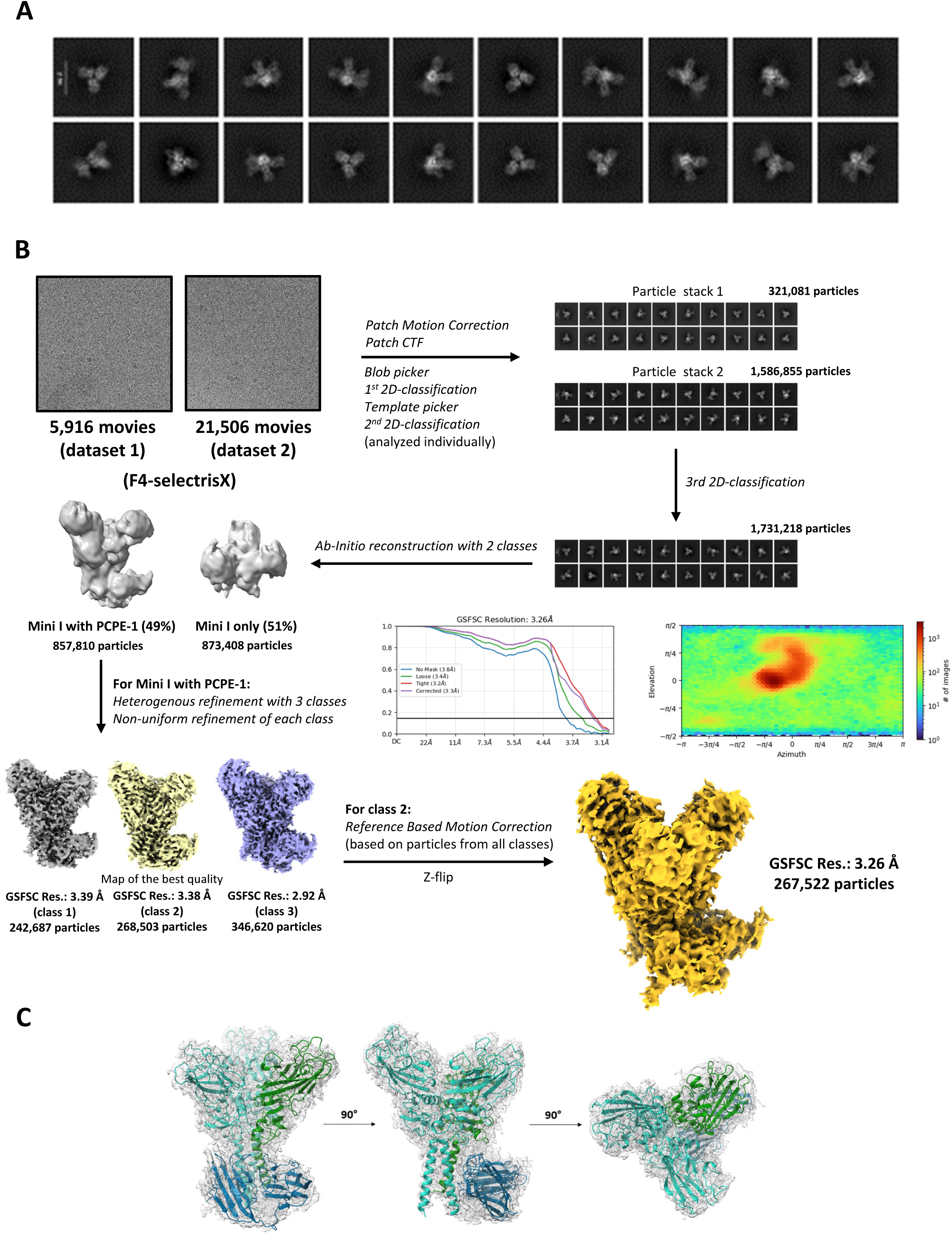
Cryo-EM data analysis workflow for the WT heteroMini-I/PCPE-1 complex. **(A)** Representative 2D class averages derived from the cryo-EM data processing workflow. **(B)** Workflow illustrating the cryo-EM data processing leading to the generation of the consensus map at 3.26 Å resolution. Fourier Shell Correlation (FSC) plots of the final map are shown (upper left of the consensus map), together with the viewing direction distribution plot (upper right). **(C)** Model-to-map fit of the refined heteroMini-I/PCPE-1 structure into the 3.26 Å cryo-EM density map.

**Supplementary Figure S6:**
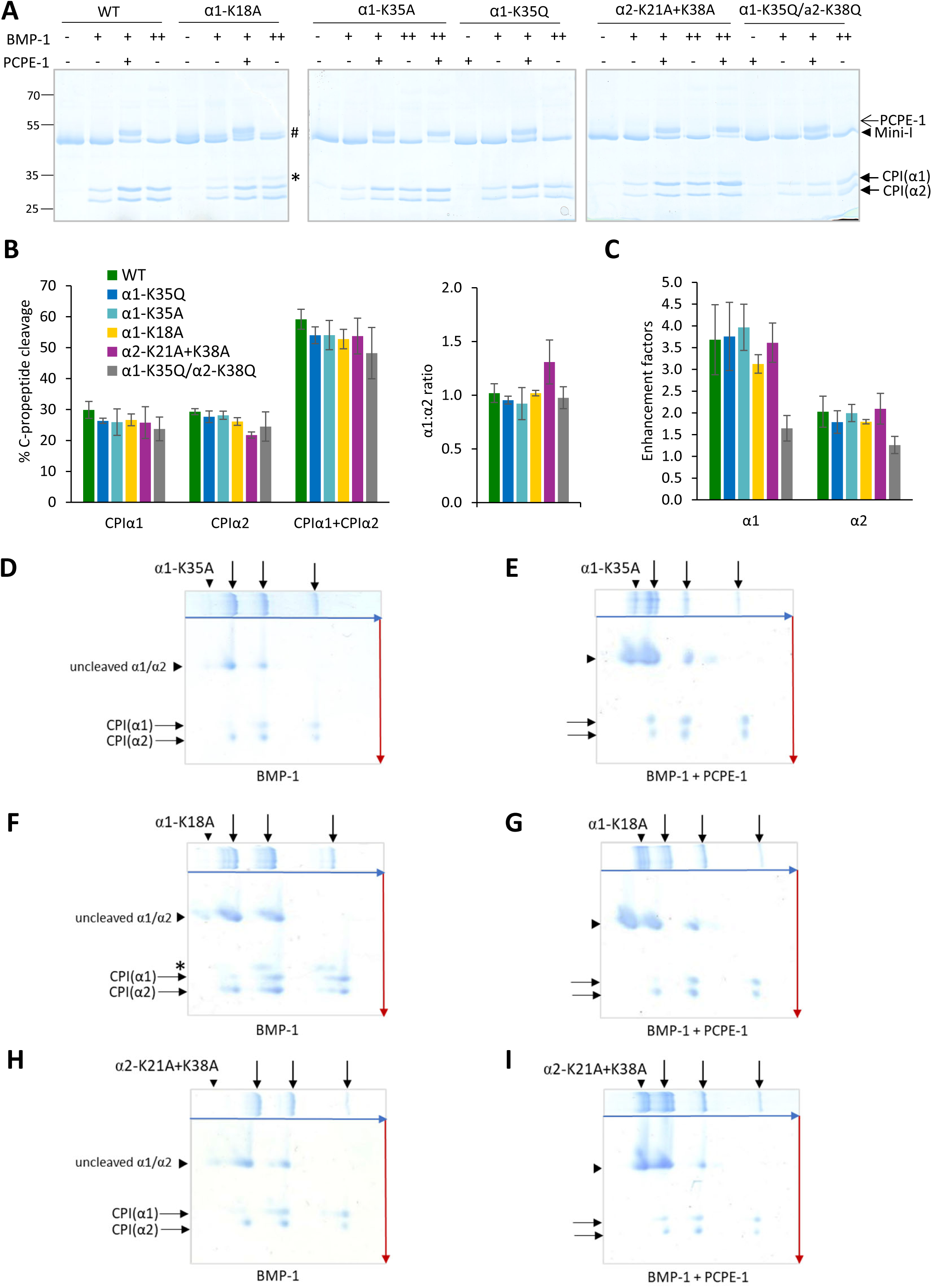
Analysis of BMP-1–mediated cleavages of heteroMini-I binding mutants. **(A)** Representative 1D gels (11 % acrylamide, reducing conditions) used to analyze BMP-1 and PCPE-1 activities on individual α chains with WT and mutant heteroMini-I (450 nM). Conditions of low (4 nM BMP-1, 0.5 h; indicated by “+”) or intermediate (20 nM BMP-1, 0.5 h; indicated by “++”) basal cleavage, with or without 450 nM PCPE-1, are shown. # and * indicates additional bands, observed in both uncleaved and cleaved α1-K18A mutant, which are not present in WT. As K18 is surrounded by several threonine and serine (Fig. S1), these products were attributed to additional O-glycosylation(s) in the absence of the bulky lysine in the mutant. These products were therefore counted as originating from the α1 chain in all quantifications. **(B)** Comparison of cleavage efficiencies and α1:α2 ratios for WT and heteroMini-I binding mutants (450 nM) in conditions of intermediate basal BMP-1 activity (cleavage efficiencies in conditions of low basal activity are shown in Fig. 4C). Cleavage assays were analyzed in 1D SDS-PAGE like in (A) followed by quantification of band intensities. Means ± SD of n = 5 (WT and α1-K35A) or n = 3 (other mutants) independent experiments. **(C)** Enhancement factors were calculated in conditions of low basal BMP-1 activity from 1D-gel densitometric analysis (same color code as in (B)). **(D-I)** Representative 2D gels showing the extent of cleavage of individual chains in α1-K35A (D, E), α1-K18A (F, G) and α2-K21A+K38A (H, I). Cleavage conditions were the same as in Fig. 2B and gels were obtained in the absence (D, F, H) and presence (E, G, I) of PCPE-1. Quantification is shown in Fig. 4C and D. * indicates the additional cleavage product of the α1 chain of α1-K18A, also seen in 1D gels (A).

**Supplementary Figure S7:**
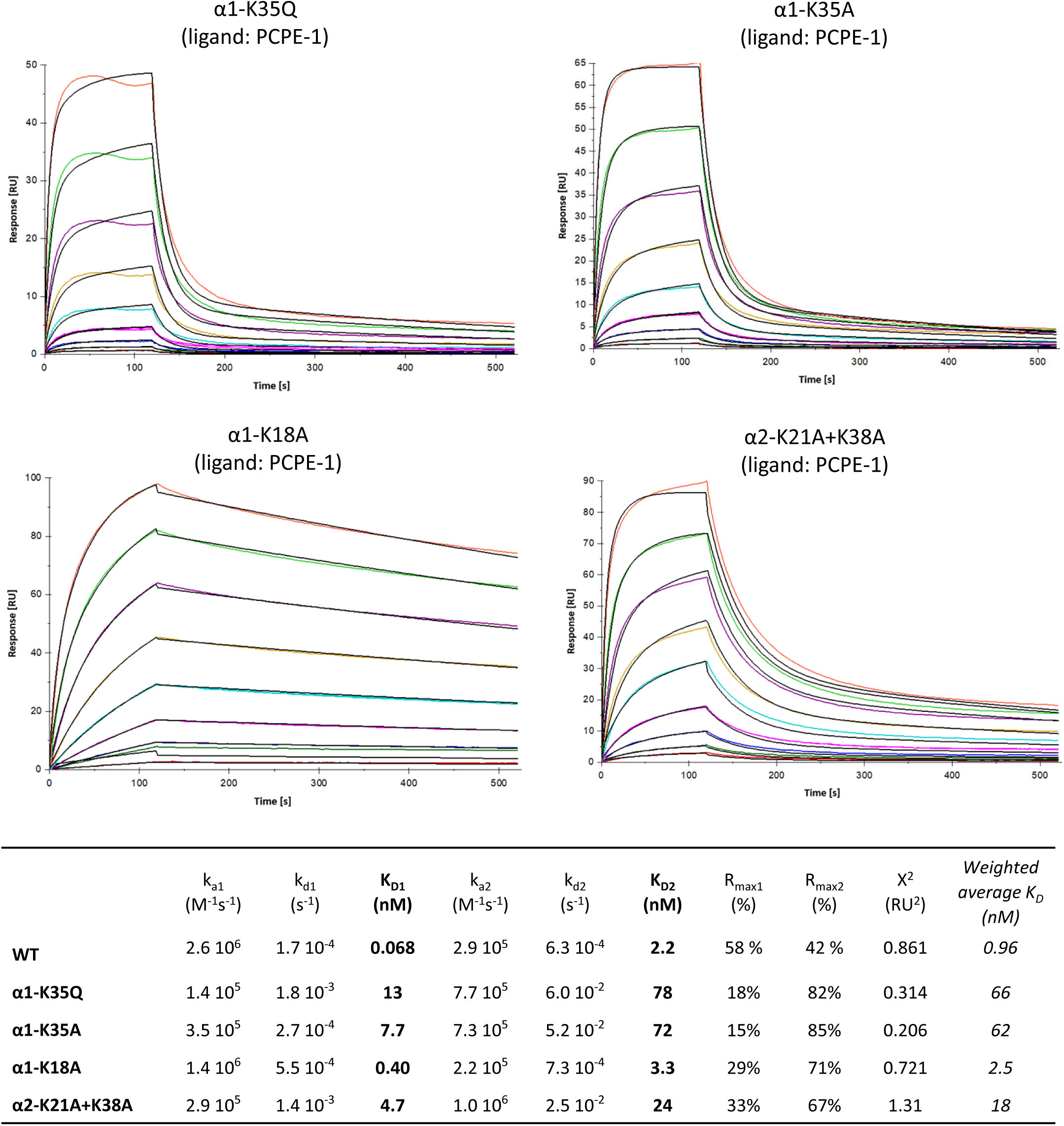
SPR sensorgrams showing the interaction of heteroMini-I binding mutants with immobilized PCPE-1 (140 RU). Increasing concentrations of the mutant proteins (0.39–100 nM; serial two-fold dilutions, including a duplicate injection at 6.25 nM) were injected, except for the α1-K35Q and α1-K35A mutants, for which the dilution series was between 200 nM and 0.78 nM. The best fits of the experimental sensorgrams obtained using the heterogeneous ligand model are shown (black curves), and the corresponding kinetic and dissociation constants are listed in the table below. For the ease of comparison, we also calculated a “weighted average K_D_” which is the mean of K_D1_ and K_D2_ weighted by the contributions of R_max1_ and R_max2_ to the total signal.

**Supplementary Figure S8:**
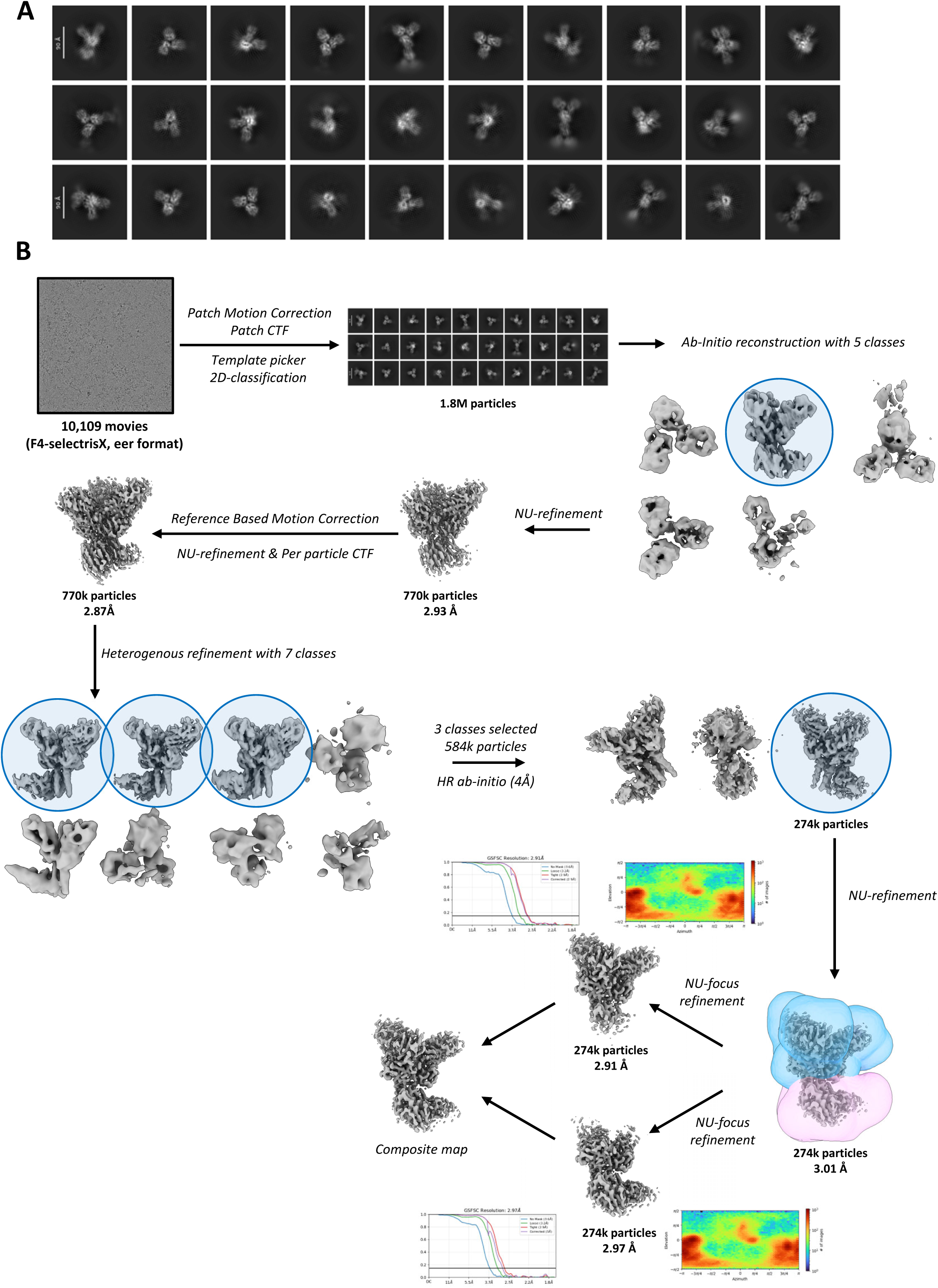
Cryo-EM data analysis workflow for the α1-K18A heteroMini-I/PCPE-1 complex. **(A)** Representative 2D class averages derived from the cryo-EM data processing workflow. **(B)** Workflow illustrating the cryo-EM data-processing procedure leading to the generation of the consensus map from focused maps at 2.91 Å and 2.97 Å resolution for the petal region and PCPE-1-binding region, respectively. Fourier shell correlation (FSC) plots of the focused maps are shown in the upper left corner of each focused map, together with the corresponding viewing-direction distribution plots in the upper right corner.

**Supplementary Figure S9:**
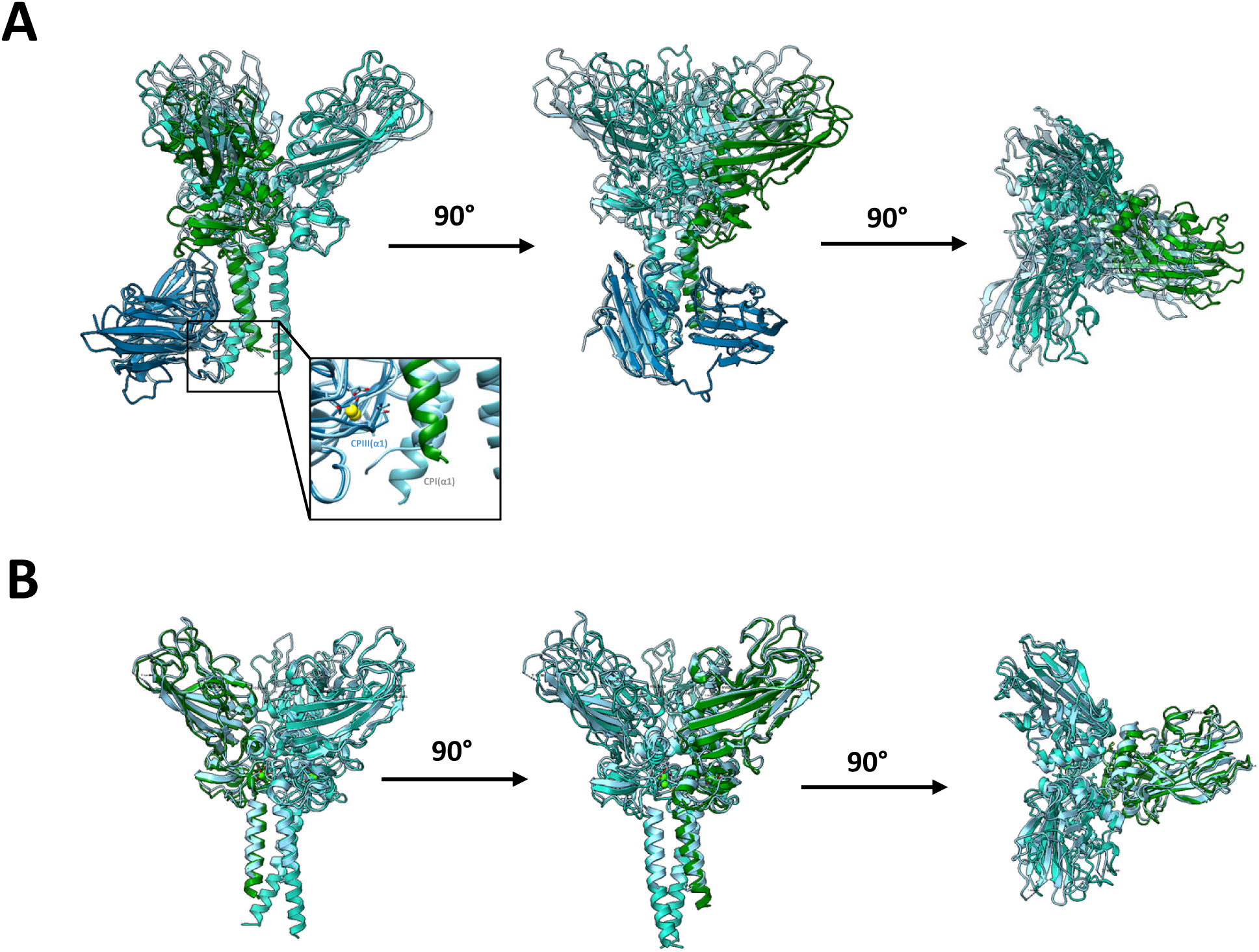
Structural superposition of the heteroMini-I/PCPE-1 model with previously reported CPIII structures. **(A)** Superposition of the WT heteroMini-I/PCPE-1 structure with the CPIII/PCPE-1 complex (PDB ID: 6FZW; blue), aligned using the coiled-coil region. The inset shows a close-up view on the disruption of chain alignment previously observed in the CPIII α1 chain binding CUB1. **(B)** Superposition of heteroMini-I, extracted from the WT heteroMini-I/PCPE-1 structure, with CPIII alone (PDB ID: 4AK3; blue), aligned using the coiled-coil region.

## Supplementary Tables

**Table S1:** Summary of data collection, image processing, and model refinement parameters.

| Data collection and processing |  |  |
| --- | --- | --- |
| **PDB entry** | 9SHT | 29OC |
| **Extended PDB entry** | pdb_00009sht | pdb_000029oc |
| **EMBD entry** | EMD-54910 | EMD-57259 (consensus map)<br>EMD-57260 (focused map 1)<br>EMD-57261 (focused map 2) |
| **Voltage (kV)** | 300 | 300 |
| **Magnification** | 165k | 165k |
| **Electron exposure (e<sup>-</sup>/Å<sup>2</sup>)** | 45 | 50 |
| **Defocus range (μm)** | -0.8 to -2.4 | -1.4 to -2.4 |
| **Pixel size (Å/pixel)** | 0.73 | 0.73 |
| **Symmetry imposed** | C1 | C1 |
| **Particle images (no.)** | 267,522 | 586,000 |
| **Map global resolution (Å)** | 3.26 | 3.01 |
| **Local resolution FSC threshold** | 0.143 | 0.143 |
| Refinement |  |  |
| **Model composition** |  |  |
| No. of protein chains | 2/1/1 | 2/1/1 |
| Non-hydrogen atoms | 7,571 | 7,570 |
| Residues | 968 | 969 |
| **Validation** |  |  |
| MolProbity score | 1.52 | 1.90 |
| Clash score | 8.84 | 7.09 |
| Rotamer outliers (%) | 0 | 1.8 |
| **R. M. D deviations** |  |  |
| Bond length (Å) | 0.004 | 0.003 |
| Bond angle (°) | 0.990 | 0.553 |
| **Ramachandran plot (%)** |  |  |
| Outliers | 0.21 | 0.10 |
| Allowed | 1.98 | 4.37 |
| Favoured | 97.81 | 95.53 |
| Rama-Z-score | 0.33 | -0.61 |
| **Model vs. data** |  |  |
| CC (mask) | 0.57 | 0.82 |
| CC (volume) | 0.56 | 0.80 |

**Table S2:** List of interactions contributing to the stabilization of the interfaces in α1-K18A heterotrimeric C-propeptide I. These interactions were identified using the PDBePISA server in structure 29OC (α1-K18A heteroMini-I in complex with PCPE-1, as described in Fig. 5) and categorized into hydrogen bonds, salt bridges, and disulfide bonds, along with the corresponding ΔG values and interface areas. The values in brackets were calculated including the interchain disulfide bonds which were not modeled in the structure.

| Interface | Interface area [Å<sup>2</sup>] | $\Delta G$ [kcal/mol] | Hydrogen bonds | Salt bridges | Interchain disulfide bonds |
| --- | --- | --- | --- | --- | --- |
| **α1-α2** | 941.5 | -13.7 | Cys64-Asp46<br>Cys64-Ser50<br>Asn61-Arg42 | Arg9-Asp13 | - |
| **α2-α1** | 910.8<br>(921.5) | -16.2<br>(-17.7) | Thr29-Arg28<br>Gln26-Arg28<br>Thr133-Arg42<br>Lys12-Asp10<br>Lys129-Glu143 | Lys129-Glu143<br>Lys12-Asp10 | Not visible<br>(Cys67-Cys47) |
| **α1-α1** | 826.4<br>(841.6) | -14.3<br>(-16.4) | Arg39-Asn61<br>Arg42-Asp67<br>Arg42-Asp129<br>Asp10-Arg9<br>Asp43-Cys64 | Arg42-Asp67<br>Arg42-Asp129 | Not visible<br>(Cys64-Cys47) |

